# Regulation of TRIB1 abundance in hepatoma models

**DOI:** 10.1101/2022.07.07.499132

**Authors:** Sébastien Soubeyrand, Paulina Lau, Ruth McPherson

## Abstract

Tribbles related homolog 1 (TRIB1) contributes to lipid and glucose homeostasis by facilitating the degradation of cognate cargos by the proteasome. We previously reported that TRIB1 was unstable in non-hepatic cellular models. Moreover, inclusion of proteasome inhibitors failed to prevent TRIB1 loss, consistent with the involvement of proteasome independent degradative processes. In view of the key role of TRIB1 in liver function, we continue our exploration of TRIB1 regulation pathways in two commonly used human hepatocyte models, HuH-7 and HepG2 cells. Proteasome inhibitors potently upregulated both endogenous and recombinant TRIB1 mRNA and protein levels. Increased transcript abundance was independent of MAPK activation while ER stress was a relatively mild inducer. Despite increasing TRIB1 protein abundance and stabilizing bulk ubiquitination, proteasome inhibition failed to stabilize TRIB1, pointing to the predominance of proteasome independent protein degradation processes controlling TRIB1 protein abundance in hepatomas. Proteasome inhibition via downregulation of its PSMB3 regulatory subunit, in contrast to its chemical inhibition, had minimal impact on TRIB1 levels. Moreover, immunoprecipitation experiments showed no evidence of TRIB1 ubiquitination. Cytoplasmic retained TRIB1 was unstable, indicating that TRIB1 lability is regulated prior to its nuclear import. Substitution of the TRIB1 PEST-like region with a GST helical region or N-terminal deletions failed to fully stabilize TRIB1. Finally, inclusion of protease or autophagy inhibitors *in vivo* did not rescue TRIB1 stability. This work excludes proteasome-mediated degradation as a significant contributor to TRIB1 instability and identifies transcriptional regulation as a prominent mechanism regulating TRIB1 abundance in liver models in response to proteasome inhibition.

## Introduction

The TRIBBLES family comprises 3 evolutionary divergent forms in higher eukaryotes: TRIB1, TRIB2 and TRIB3. A distinguishing feature is their ability to promote degradation of cognate targets by enforcing their interaction with specific E3 ligases. This functional network plays an important role in normal liver function by regulating lipid metabolism but is also implicated in the development and progression of myeloblastic leukemia [1–5]. Mammalian tribbles have also been shown to regulate MAPK signaling in a coordinated fashion [4, 6]. In D. Melanogaster, where a single Tribbles is present, tribbles promotes rapid degradation of slbo turnover and ensures proper oogenesis [7]. In addition, Tribbles inhibits cell cycle progression by facilitating String and Twine proteasomal degradation [8]. Recently, the *TRIB1* gene locus has renewed interest through its association with coronary artery disease (CAD). Although it is still unclear whether *TRIB1* proper mediates the genetic association with CAD, compelling evidence points to its role in lipid metabolism [5, 9].

Tribbles are pseudokinases, namely proteins that lack phosphorylation activity but leverage structural motifs shared by kinases [10]. Over the years, mammalian *tribbles* have been shown to interact with several proteins. The best characterized interaction is with the E3 ligase COnstitutively Photomorphogenic 1 (COP1). COP1 is a RING finger E3 ligase, which in complex with E1 and E2 enzymes can catalyze ubiquitination of target proteins, in some cases with the assistance of additional adaptor proteins. By binding and recruiting CEBP to COP1, TRIB1 and TRIB2 enable degradation of CEBPs [11–15]. While all 3 tribbles can constitutively interact with COP1 via their C-termini Val-Pro containing motif, only TRIB1 and TRIB2 target CEBP for degradation, indicating that interaction might not be sufficient for degradation; by contrast, all three can facilitate COP1 mediated ACC1 degradation when overexpressed [1].

The fate of TRIB1 as an adaptor protein is unclear. E3 adaptor proteins, including TRIB1, tend to be unstable [16]. Limited evidence however suggests that it may not be degraded through the process of cargo degradation [17]. For TRIB3, COP1 overexpression reduces CEBPA without affecting TRIB3 levels, suggesting that TRIB3 is not degraded via COP1 [1]. Thus, TRIB1 may not be directly degraded by the ubiquitin proteasome system (UPS). However the Val-Pro interface, mediating binding of both adaptor proteins and targets to COP1, also plays a key role in earmarking degradation targets (e.g. Jun, ATGL, ETS) [18, 19].

Regulated protein degradation involves three major pathways. Removal of ubiquitinated proteins typically occurs via the 26S, a large protein complex exhibiting 3 distinct protease activities [20–22]. Several proteasome inhibitors (PIs) have been developed over the years, some of which have clinical implications [23, 24]. Chief among them is Velcade/Bortezomib (BTZ), used in the treatment of multiple myeloma. Relevant to liver disease, BTZ therapy is sporadically linked to liver injury and has been shown to exhibit hepatotoxicity in mouse models [25].

In addition to proteasome-mediated degradation, lysosomes and autophagosomes also contribute to the removal of larger cargos via lysosomal proteases [21, 26]. All 3 pathways share some dependence on ubiquitination and are likely functionally interconnected, particularly under stress [27]. For instance, interfering with proteasome function can induce autophagy to help tackle the accumulation of unwanted protein conjugates and contribute to homeostasis.

We previously demonstrated that TRIB1 regulation in non-liver models occurred at multiple levels, involving proteasome dependent and independent processes at transcriptional and post-transcriptional levels [28]. An intriguing finding was that whereas drug mediated proteasome inhibition was sufficient to increase TRIB1 levels, associated with increased transcription, proteasome inhibition was insufficient to prevent loss of TRIB1 after CHX treatment, hinting to alternate pathways that could degrade TRIB1. The current study examines the regulatory processes controlling TRIB1 abundance in greater detail, leveraging two commonly used human hepatocyte cell models.

## Materials and Methods

### Cell culture and treatments

HuH-7 (Japanese Collection of Research Bioresources Cell Bank) and HepG2 (ATCC) cells were maintained in physiological (5 mM) glucose DMEM media supplemented to 10% FBS and containing antimycin A and antibacterial supplements (Gibco). Drugs are listed in the **Supplemental Material** section. Stable cell pools were obtained by 3 ug/ml puromycin selection following viral transduction. Vehicle (DMSO or PBS) was added at 0.1% or less, final concentration.

### Western blotting

Protein lysates (approximately 30 µl per lane) were resolved on 1.5 mm 8 % SDS-PAGE gels unless mentioned otherwise and transferred to nitrocellulose membranes. Membranes were blocked in Intercept buffer (LI-COR, 30 min), rinse briefly in PBS/0.1% Tween (PBS/T) and incubation with the primary was typically performed in PBS/T for 16 h. Secondary antibodies were from Li-Cor (680 and 800) and were used as 1:20,000 dilutions. Antibody incubations were followed by one PBS/T and three PBS rinses. All antibodies used are listed in the **Supplemental Material** section. TRIB1 is expressed at very low levels and was undetectable in naïve HuH-7 and HepG2 using 2 distinct Abs under basal conditions; lack of TRIB1 signal was ascertained using orthogonal validation using antisense oligonucleotides and lack of responsiveness to actinomycin D. Different lots of TRIB1 antibodies raised in rabbits (TRIB1r; targeting the N-terminal region) were inconsistent in sensitivity but yielded comparable background signals. The goat TRIB1 (TRIB1g) was more consistent across lots but exhibited slightly lower sensitivity. Analyses were performed with either or both Ab as limited supplies permitted. Endogenous and exogenous wild-type TRIB1 were detectable as a tight band triplet with a prominent central band. Note that both Abs revealed the presence of an overlapping non-specific band (resistant to TRIB1 siRNA and ACTD).

### Immunoprecipitation

Whole cell lysates of HepG2-T1 (stably transduced with pLVXpuro-TRIB1) cells were obtained by lysing cells in PBS containing 0.1% Triton X-100 (PBS/TX) as well as protease and protein inhibitor cocktails (Roche) for 5 min on ice. Cleared lysates (13,000 xg for 2 min @ 4 C) were adjusted to 1 mg/ml with PBS/TX buffer and antibodies were added at 2 ug/mg lysate for 30 min, followed by the addition of protein A/G beads (10 ul suspension; PureProteome, EMD Millipore) for an additional 5 h on an orbital plate. Samples were then washed 4 X with 0.5 ml of IP buffer, heat denatured at 95 °C under reducing conditions (5 min) and analyzed by Western blot.

### Real-Time RNA quantification (RT-qPCR)

RNA was extracted from culture plates using Tri Reagent (Roche) and isolated using Direct-Zol RNA miniprep kits (Zymo Research). RNA was reverse transcribed using a mixture of oligo dT and random hexamer primers (Roche). Quantification was performed on a Light Cycler 480 using SYBR Green (Roche). Target of interest values were expressed relative to the corresponding peptdidyl Peptidylprolyl Isomerase A (PPIA) values using the 2−ΔΔCt method. Oligonucleotides are listed in the **Supplemental Material** section.

### Plasmid and lentiviral constructs

TRIB1NES, wherein the bipartite TRIB1 nuclear localization sequence (AA 31-51) was substituted for a PKIA nuclear export signal as described previously, was subcloned into PLVX-puro [29]. For the PEST substituted construct, a synthetic fragment containing the GST helical region, consisting of 2 identical helices, was used to replace the PEST region (AA 53-88) of TRIB1 (see **Supplemental Material** section for sequence). N-terminal truncated TRIB1 expression constructs (Δ2-51 and Δ2-91) were obtained by sub-cloning TRIB1 from the corresponding pCFP-TRIB1 construct to pLVX via BamHI transfer. Lentiviral constructs were packaged in 293FT cells using co-transfection of pLVX constructs with with psPAX2 and pMD2.G (Addgene). Virus titers were estimated using puromycin resistance in HEK-293T cells; an equivalent of 3 MOI was used per infection.

### Microscopy

Cells were seeded on glass coverslips and fixed with 4% PFA in PBS for 15 min and subsequently permeabilized/fixed for 20 min in PBS containing 0.1% Triton X-100 and 5% FBS. Detection was performed by incubating with primary antibodies at 1:500 dilutions in PBS and secondaries (Jackson laboratories) at 1:2000 for 1 h in PBS/0.1% Triton. Four 1-minute rinses in PBS were performed after each incubation. Cells were mounted in Fluorescent Mounting Medium (DakoCytomation) supplemented with Hoechst 33342. Microscopy was performed using a 63 X Oil Immersion lens on a Zeiss Elyra microscope.

## Results

### TRIB1 abundance in hepatocyte models is upregulated by proteasome inhibitors

In liver models, the PI bortezomib (BTZ) increased recombinant TRIB1 and unexpectedly revealed a weak TRIB1 signal in vector transduced HepG2 (Fig 1A). The presence of multiple novel bands was confirmed in naïve HepG2 but not in HuH-7 (Fig 1B). Moreover, as previously observed in non-liver models, TRIB1 was unstable, as revealed by cycloheximide (CHX) treatment, resulting in the disappearance of the TRIB1 signal within 5 h of the translation inhibitor (Fig 1B). Increased TRIB1 was accompanied by a 3 to 14-fold TRIB1 transcript increase in HepG2 and HuH-7 cells, respectively and was observed with other, mechanistically, and structurally distinct, inhibitors Epoxomycine (EPOXO), Lactacystin (LACTA) (S1 Fig). With prolonged incubation, TRIB1 was detected in both parental HepG2 and HuH-7 cells, indicating that TRIB1 accumulates over time (Fig 2). Immunofluorescence revealed a weak TRIB1 reactive signal that was enriched in the nucleus following BTZ treatment, detectable after as early as 5 h in HepG2 cells or after longer incubation in HuH-7(Fig 3, S2 Fig, S3 Fig).

**Fig 1.**
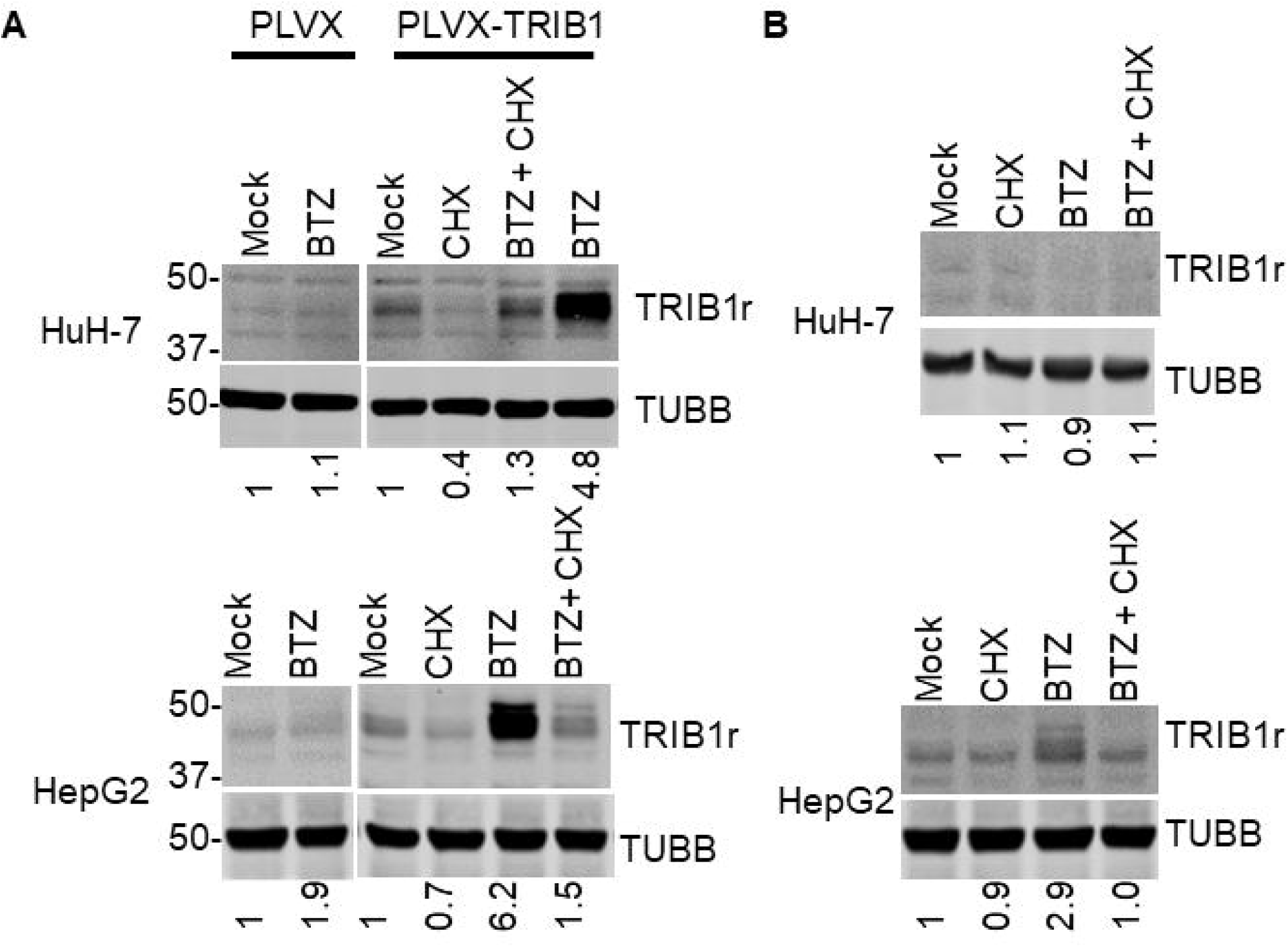
Overexpressed TRIB1 and endogenous TRIB1 are sensitive to CHX and proteasome inhibition in HuH-7 and HepG2 cell models. A, PLVX and PLVX-TRIB1 stables were exposed to 5 h BTZ treatment in presence of CHX or vehicle (1% DMSO). B, as in A, but parental HepG2 and HuH-7 were treated as indicated. Data representative of 3 experiments. Quantifications (relative to TUBB) normalized to Mock are shown.

**Fig 2.**
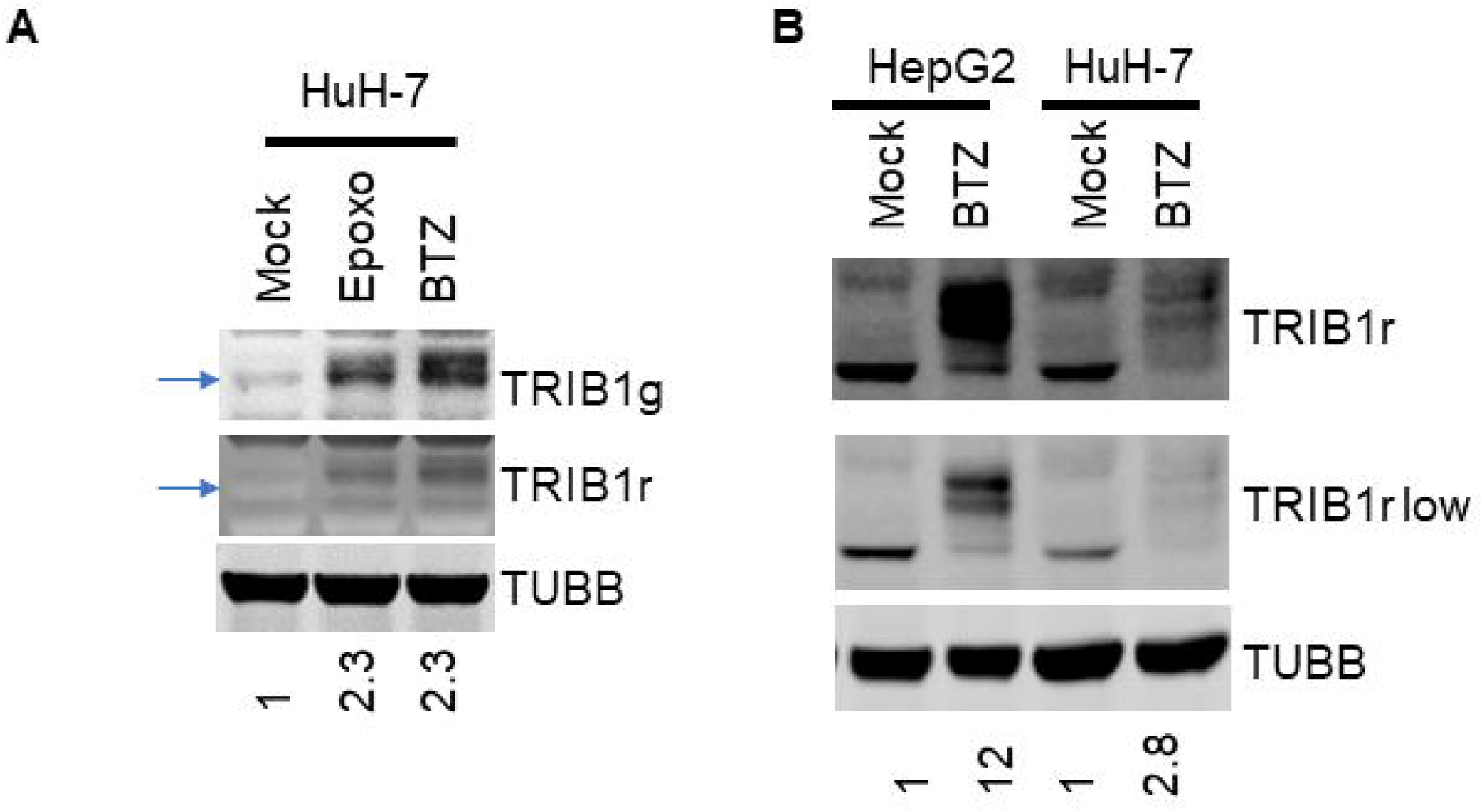
Prolonged incubation with proteasome inhibitors reveals endogenous TRIB1 in HepG2 and HuH-7 cells. A, TRIB1 in HuH-7 is increased in response to a 18 h treatment with 2 µM epoxomicin (Epoxo) or 5 µM BTZ. Arrow indicates equivalent position on both panels. Detection was performed either with goat or rabbit raised Abs (in distinct channels), as indicated. B, Relative abundance in HepG2 and HuH-7 cells following 18 h incubations with BTZ. Two different intensities are shown. Data are representative of 3-5 experiments. Quantification is expressed relative toTUBB and normalized to the Mock sample; for TRIB1 quantification, the TRIB1r signal was used.

**Fig 3.**
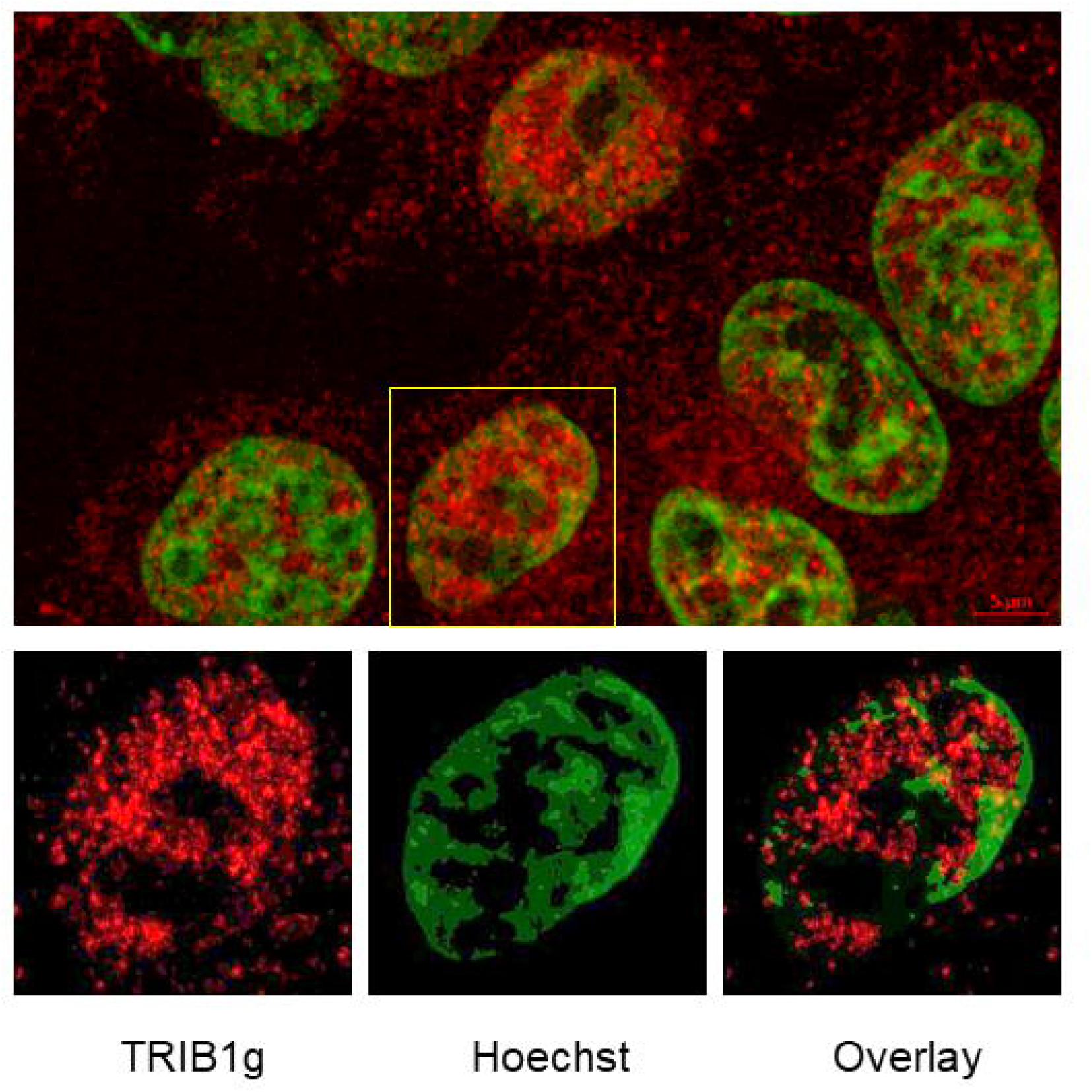
Endogenous TRIB1 signal in HepG2 cells treated with BTZ treatment for 5 h. HepG2 cells on coverslips were treated with BTZ (5 uM) for 5 h. Detection was performed using a TRIB1 Ab (TRIB1g; red) and Hoechst (green) signal. Top, representative overlay of signals from a group of cells. Bottom, individual channels of highlighted nucleus. Image was smoothed with a Gaussian filter (default parameter settings ZEN 3.3)

### Increased transcript abundance is the result of increased transcription

Increased TRIB1 abundance could reflect increased transcript output or increased stabilization. The role of transcription in regulating TRIB1 abundance in response to BTZ was examined by blocking *de novo* POLII transcription with ACTD. Comparable fold reductions in TRIB1 mRNA were observed upon treating control and BTZ treated cells with ACTD, suggesting that ongoing transcription is required (Fig 4A). Moreover, ACTD treatment was sufficient to prevent BTZ mediated protein accumulation (Fig 4B).

**Fig 4.**
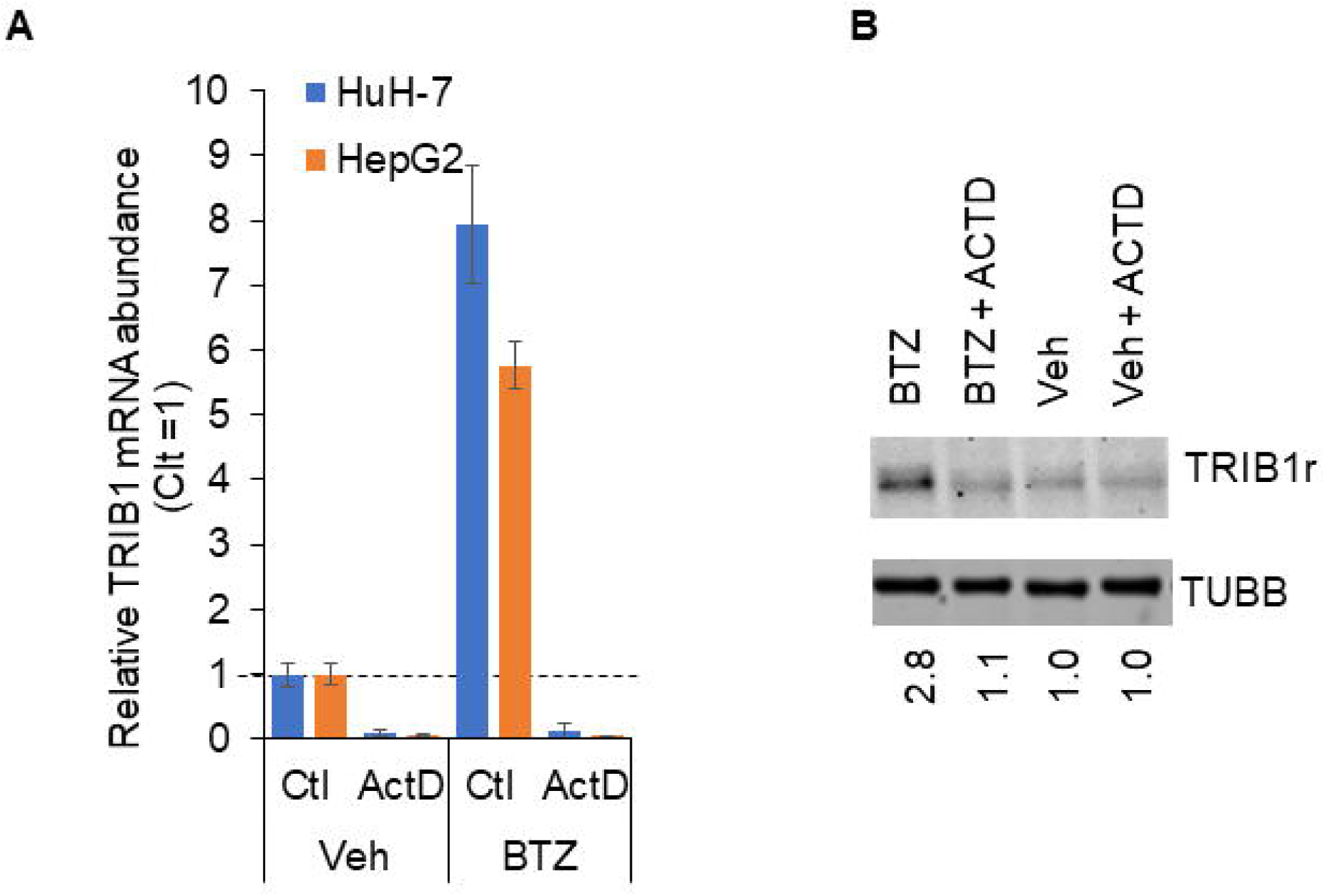
Transcription inhibition blocks TRIB1 upregulation in response to 5 h BTZ treatment. HuH-7 and HepG2 cells were treated with BTZ or vehicle (Veh), with and without (Ctl) a preincubation with ACTD for 5 min. A, mRNA abundance in HuH-7 and HepG2 cells. B, Western blot performed in HepG2 cells; endogenous protein signal was undetectable in HuH-7 after 5 h. Within each cell type, differences between Ctl and ACTD were highly significant (p <0.001, one-way repeated measures ANOVA).

### Increased TRIB1 mRNA abundance in response to proteasome inhibitors is independent of MAPK activation

MAP kinases (p38, JNK and ERK1/2) are major regulators of cell proliferation and stress response and may contribute to increased TRIB1 transcription in response to BTZ induced proteotoxic stress. We previously demonstrated the essential role of ERK1/2 in promoting TRIB1 transcription following mitochondrial stress in HepG2 cells [30]. The contribution of ERK1/2 was tested by blocking of ERK1/2 activity with PD98059; this however had no effect on the BTZ mediated increase in TRIB1 (Fig 5). Proteasome inhibition also stabilizes and activates c-Jun via JNK; Jun is a major component of AP-1 complex, which occupies the TRIB1 promoter in naïve HepG2 cells and liver according to ENCODE data [31–33]. As for p38, it can be activated by proteasome inhibition and has the potential to promote AP-1 function [34, 35]. As observed for ERK inhibition, inclusion of JNK and p38 inhibitors failed to block the BTZ mediated increase in both HuH-7 and HepG2 cells. Finally combining all 3 had minimal impact on TRIB1 upregulation by BTZ. Thus increased TRIB1 mRNA abundance does not require the contribution of MAPKs.

**Fig 5.**
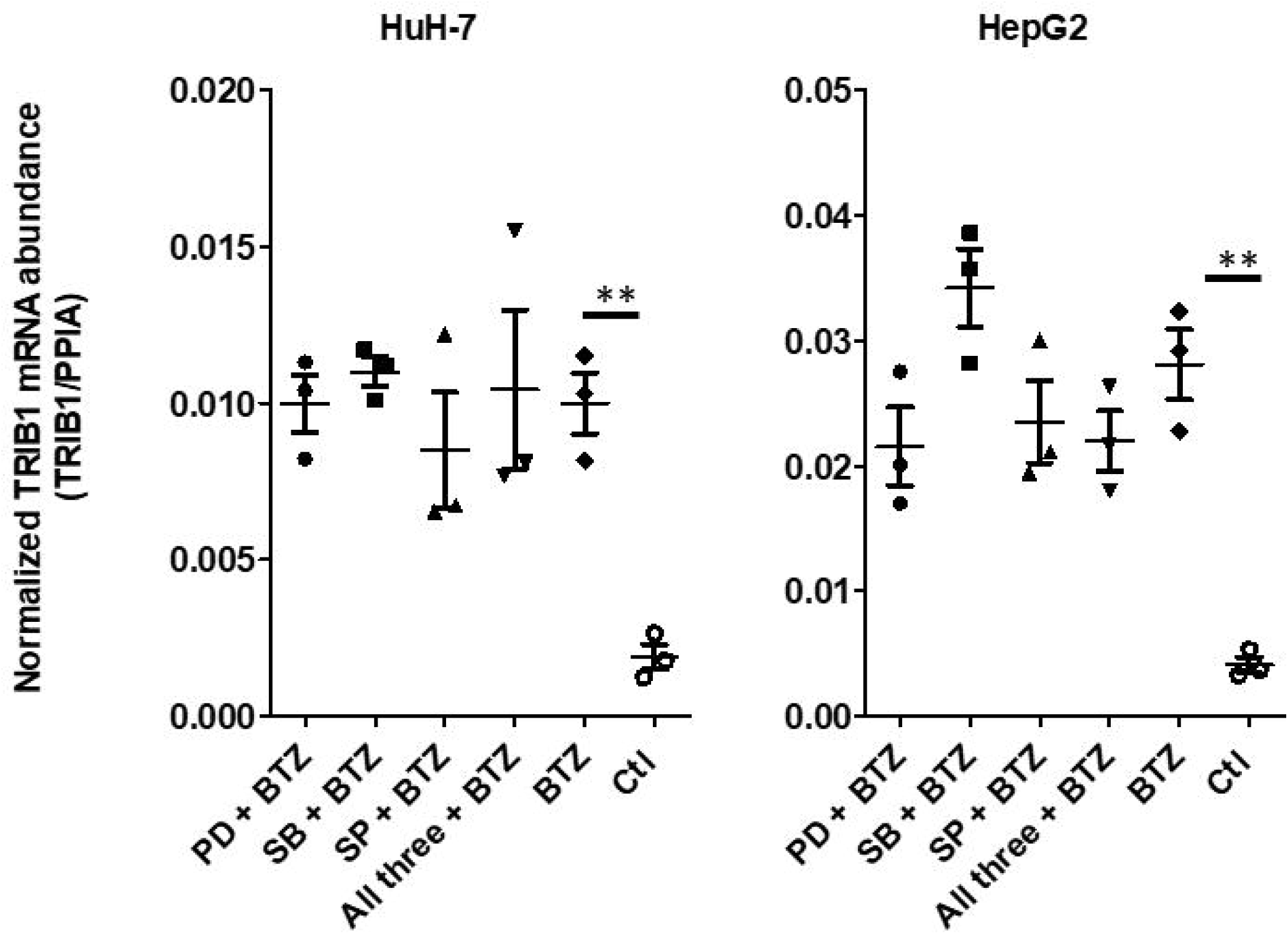
BTZ mediated increased mRNA abundance is independent of MAPK activity. HuH-7 and HepG2 cells were treated with BTZ for 5 h in the presence of MAPK inhibitors (PD: PD98059 (ERK1/2), SB: SB203580 (P38), SP: SP600125 (JNK)), either singly or in combination, as indicated; Ctl cells were treated with vehicle only (1% DMSO). Individual biological replicates are shown (± SEM). TRIB1 expression is normalized to PPIA levels. Differences between [MAPKi + BTZ] treatments and BTZ were not statistically significant (one-way repeated measures ANOVA with Dunn’s post-hoc test vs Ctl).

### TRIB1 levels are weakly responsive to ER stress

Proteasome inhibition, by interfering with protein turnover of misfolded proteins targeted for secretion, has the potential to induce ER stress and an unfolded protein response (UPR). Interestingly TRIB3 has been shown to be highly responsive to ER stress induction via tunicamycin in HepG2 cells [36]. Indeed, BTZ treatment elicited a stress response as assessed by increased CHOP and BiP in both HCC cell lines (S4 Fig). To examine the potential of ER stress in controlling TRIB1 levels, cells were treated with tunicamycin, a prototypical ER stress inducer. A 5 h (or 22 h) tunicamycin increased TRIB1 abundance modestly (1.1-2-fold); by comparison, tunicamycin treatment resulted in a 5 to 20-fold BiP and CHOP increase in both hepatoma models (Fig 6). Together, these findings indicate that ER stress occurs in response to BTZ in hepatoma models but that it is insufficient to singly account for the 4 to 15-fold increased TRIB1 mRNA levels elicited by PI treatments in HepG2 and HuH-7.

**Fig 6.**
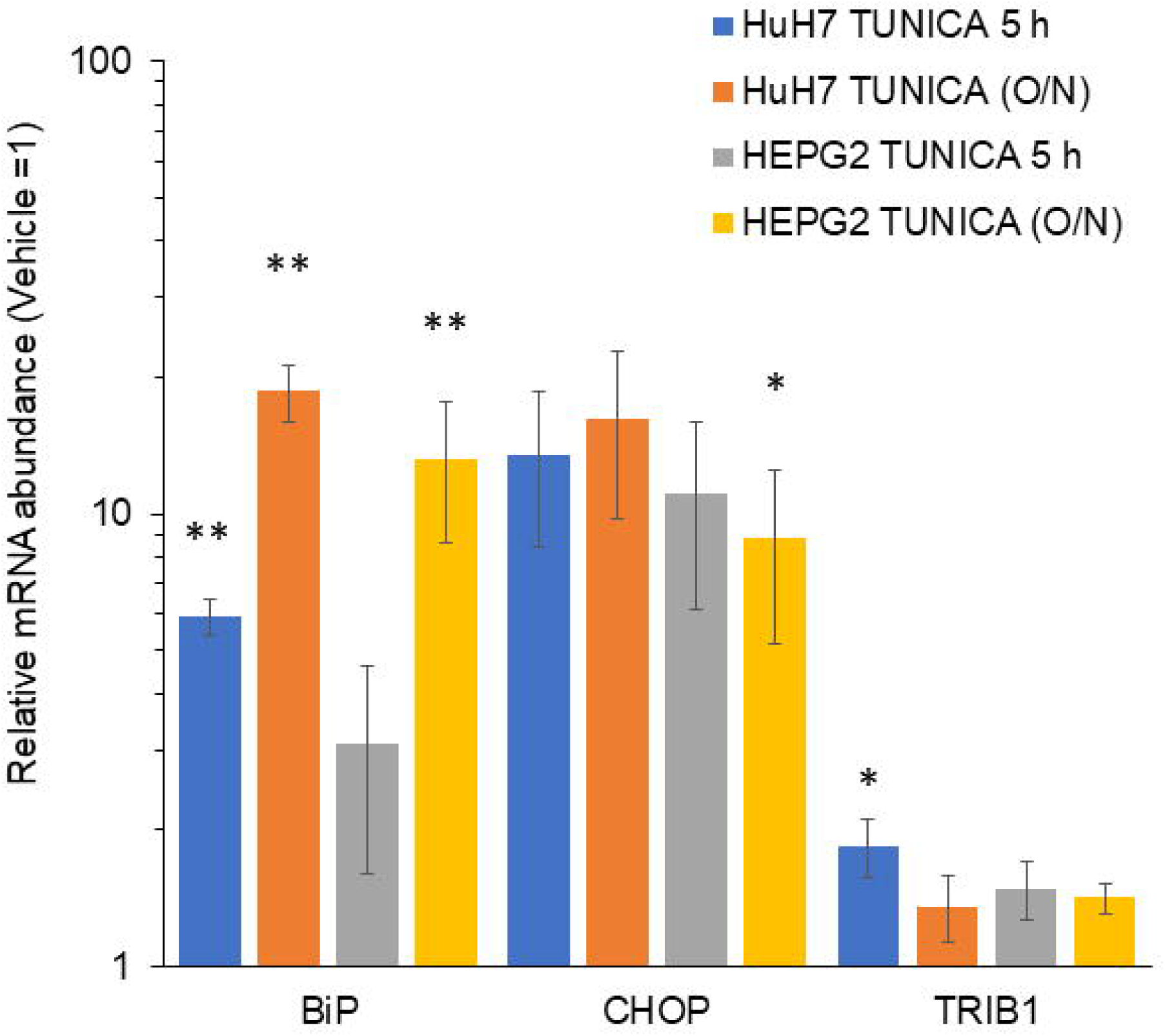
ER stress is a weak TRIB1 inducer. HuH-7 and HepG2 cells were treated with Tunicamycin (12 µM) for either 5 or 18 h (O/N). Bars represent the means +/- S/D. *, p<0.05, ** p< 0.01 from Vehicle. Paired Student t-test, Tunica vs vehicle.

### Upregulation of TRIB1 by proteasome inhibitors is rapid and is dependent on increased transcription

A time course was performed to better characterize the upregulation at the protein level. To visualize TRIB1 at early time points, when endogenous protein abundance is too low for detection, the assay was performed on recombinant TRIB1. To address possible off-target BTZ effects on non-proteasome proteases [37], lactacystin (LACTA), a structurally and mechanistically distinct inhibitor was also used [23]. Lactacystin was included at a concentration previously shown to inhibit ∼70% of the proteasome chymotryptic activity *in situ* [38]. Rapid TRIB1 upregulation was detectable within 1 h of treatment using both BTZ and LACTA and increased over 5 h (Fig 7). As observed above with the endogenous transcript, PIs potently upregulated recombinant TRIB1 mRNA (S5 Fig).

**Fig 7.**
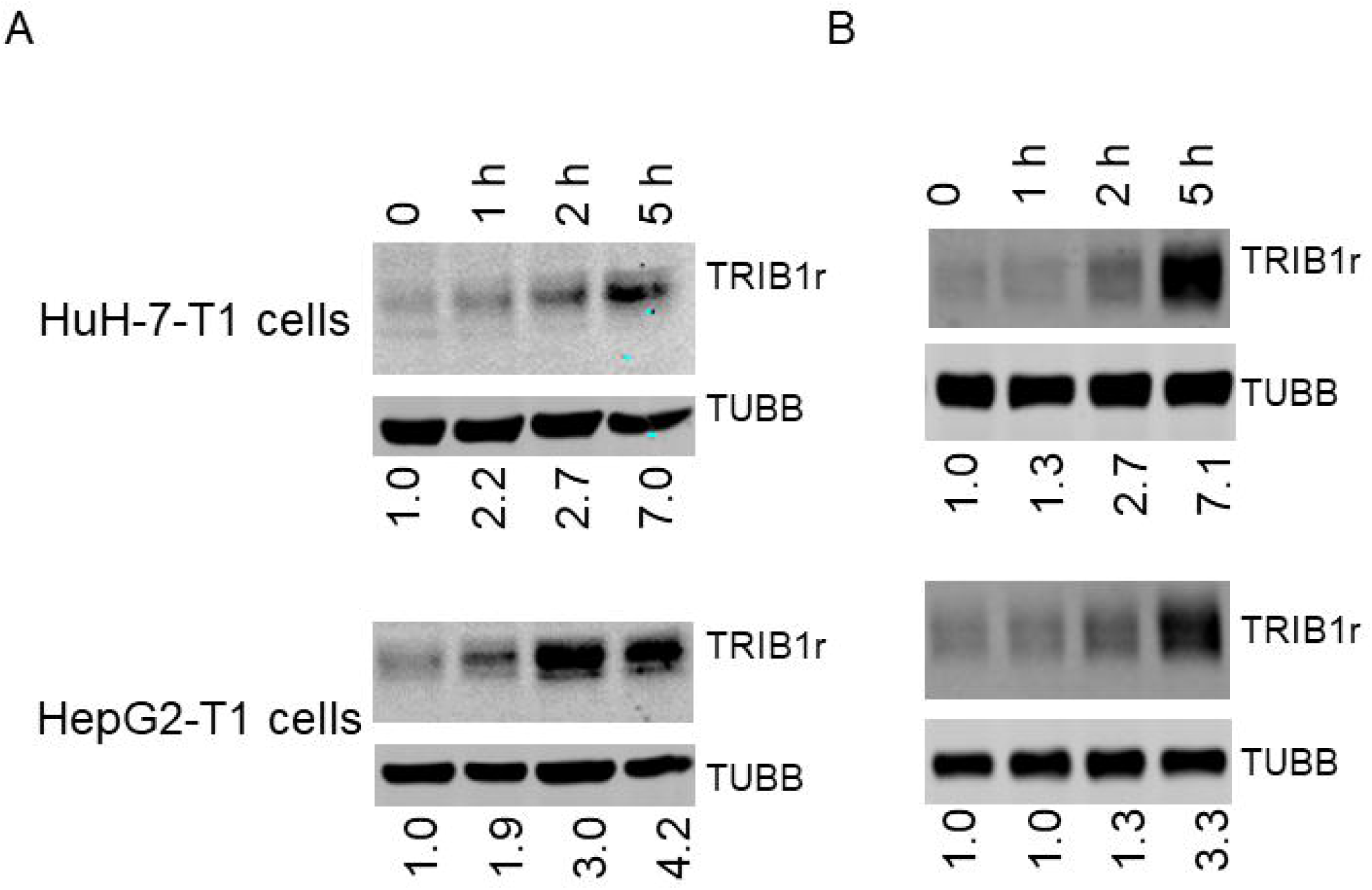
Upregulation of recTRIB1 by proteasome inhibitors is rapid. Western blot analyses of time course experiments. Stable pools of HuH-7-T1 and HepG2-T1 cells were treated for 1 to 5 h with BTZ (5 µM) (A) or LACTA 10 µ M (B) for the indicated time. Samples were resolved by Western blot and analyzed with the indicated antibodies.

Increased protein abundance could be the result of increased transcript reduced protein degradation, increased transcription, increased translation, or a combination thereof. Upregulation was blocked by ACTD, as noted earlier with the endogenous protein, indicating that increased recombinant TRIB1 protein abundance requires transcription (Fig 8).

**Fig 8.**
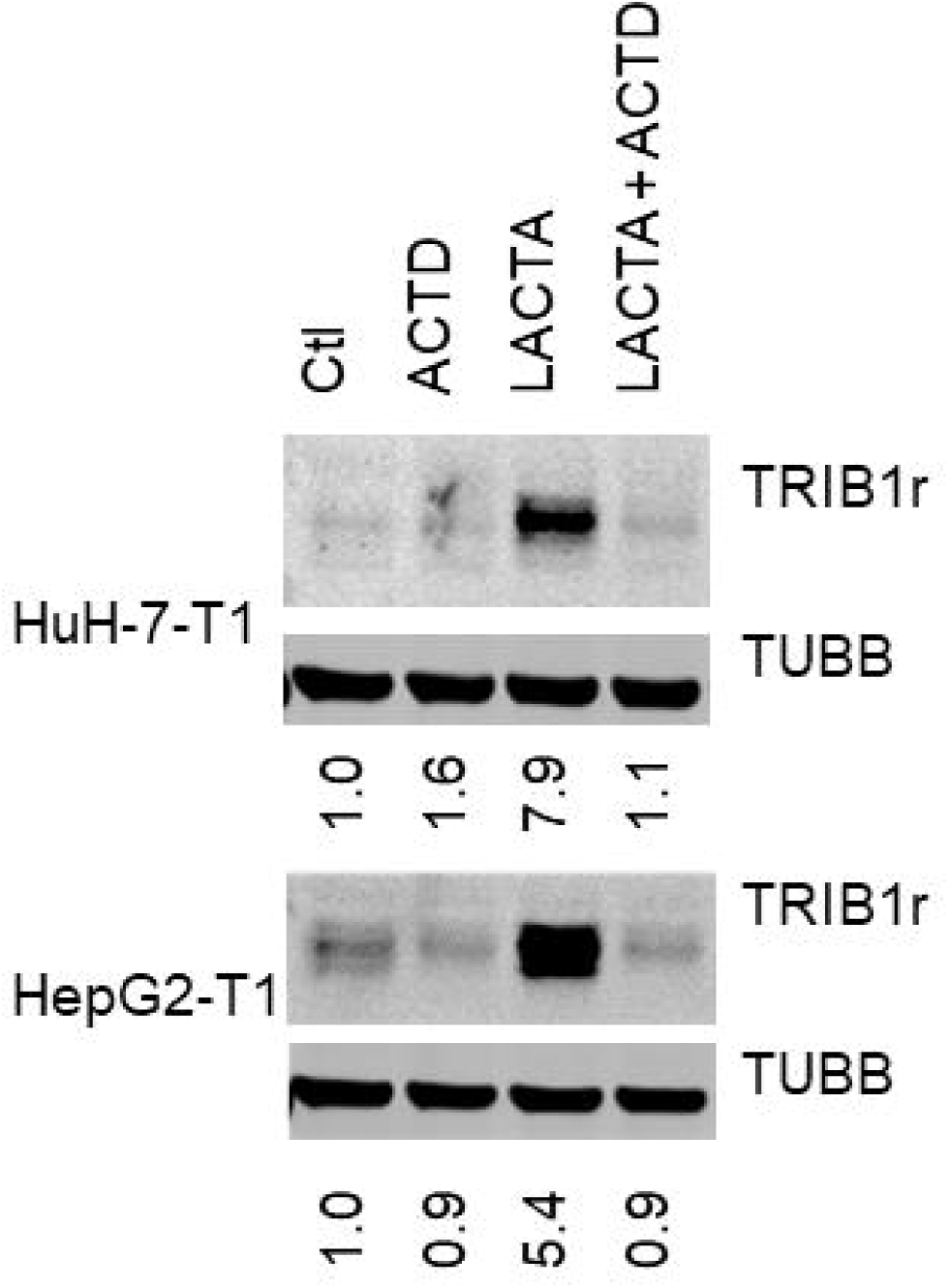
Increased recTRIB1 following proteasome inhibition is blocked by ACTD. HuH-7 and HepG2 cells stably expressing TRIB1 were incubated for 5 h with lactamycin (LACTA,10 µM) and/or Actinomycin D (ACTD, 5 µg/ml)), as indicated. Data representative of 3 experiments. Quantifications represent TUBB normalized signal intensity, expressed relative to the corresponding Ctl values.

### The proteasome plays a limited role in the control of steady state TRIB1

Evidence to date points to the major contribution of transcription in regulating TRIB1 abundance when the proteasome is blocked. These experiments however were performed in the presence of inhibitors with unavoidable off-target effects. To address these limitations, the proteasome regulatory beta 3 subunit (PSMB3) was targeted by siRNA. Reduced *PSMB3* expression resulted in increased abundance of coupled ubiquitin chains, consistent with impaired proteasome function allowing ubiquitinated proteins to accumulate. The treatment was however insufficient to reveal endogenous TRIB1 protein signal (Fig 9A). Lack of detection could be due to absent or insufficient protein upregulation. To distinguish between these 2 scenarios, proteasome inhibition via PSMB3 was repeated on cells expressing the ∼ 10-20-fold more abundant recombinant *TRIB1*. Reduced PSMB3 levels were associated with a subtle increase in TRIB1, which reached statistical significance only in HepG2-T1 (1.2-fold ± 0.07; n=3) and could be due to a correlative 1-5-2-fold increased TRIB1 mRNA (Fig 9B; S6 Fig).

**Fig 9.**
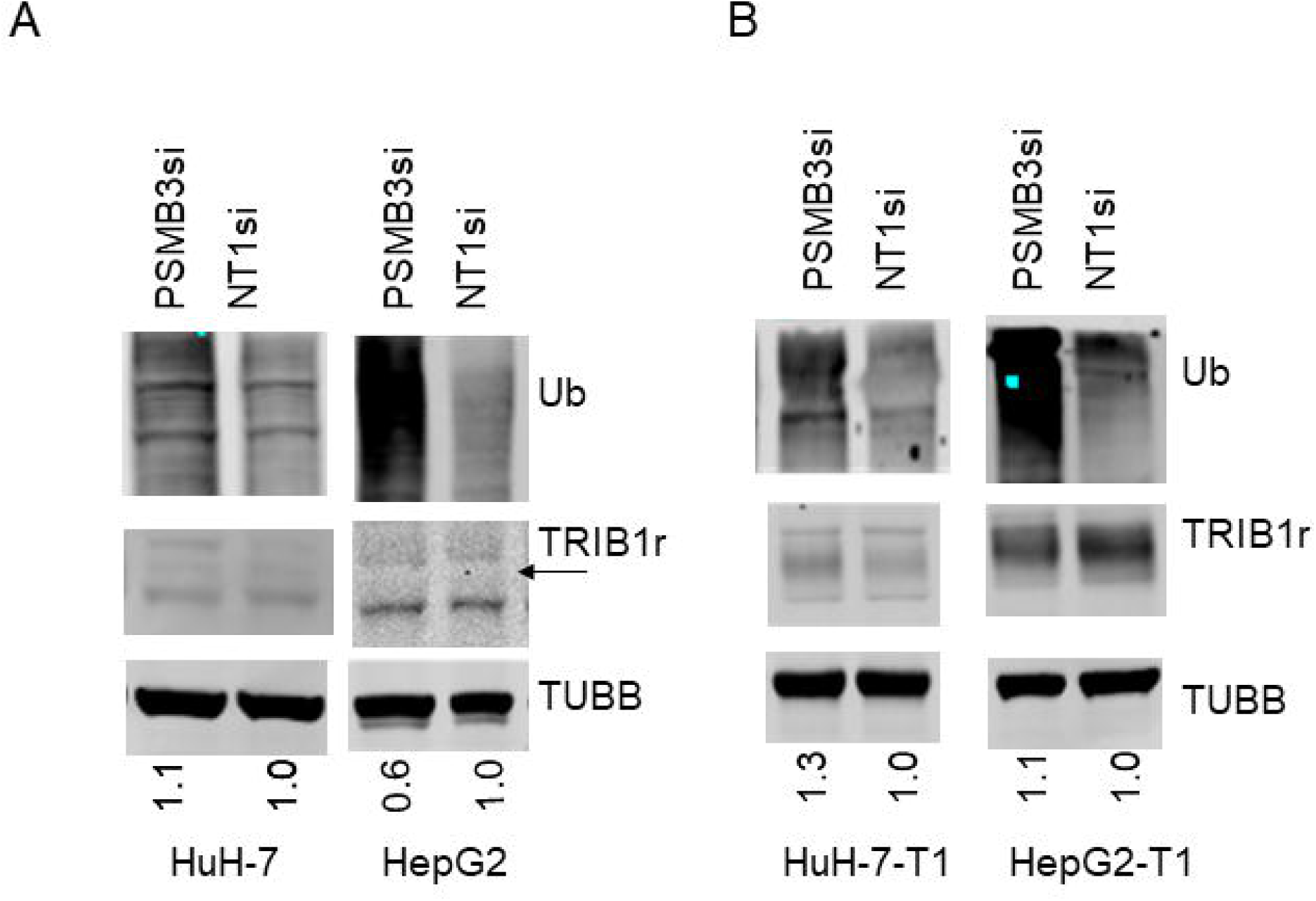
Long term proteasome suppression increases polyUb but has minimal impact on TRIB1 abundance. Western blot analyses of PSMB3 silenced samples. A, B: HuH-7 and HepG2 cells (with or without transduced TRIB1, as indicated) were treated 96 h with siRNA targeting PSMB3 or a siRNA control. C, For Ub, region spanning 100 kDa to the top of the gel is shown. Arrow points to the predicted position of TRIB1. Quantification represents the TUBB corrected TRIB1 levels, normalized to the NT control value.

### No evidence of TRIB1 ubiquitination using overexpression approaches

Evidence of TRIB1 ubiquitination was tested by performing immunoprecipitation under both basal and proteasome blocked conditions. Independent of drug treatment, no detectable ubiquitin was associated with TRIB1. With the limitation that protein ubiquitination may not always be required for proteasome-mediate degradation (e.g. [39]), these results further support the contribution of non-proteasome mediated degradation processes (Fig 10).

**Fig 10.**
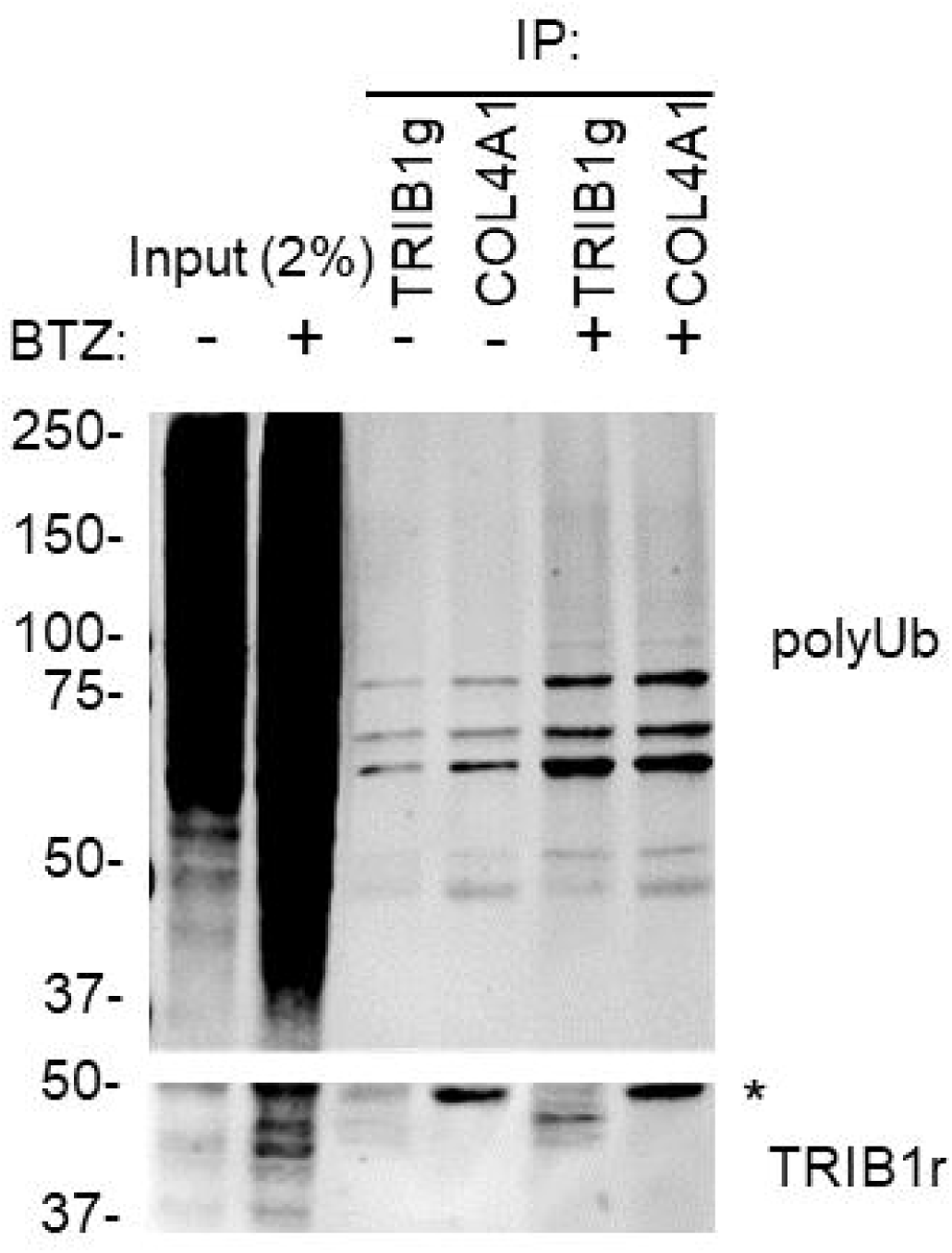
Immunoprecipitated TRIB1 shows no evidence of ubiquitination. HepG2-T1 cells, treated with either BTZ or vehicle (0.25% DMSO) for 5 h were immunoprecipitated with either TRIB1g or COL4A1 goat polyclonal antibodies and analyzed by Western blotting using rabbit anti-Ub (top) or anti-TRIB1 (bottom). Detection was sequential (Ub then TRIB1r). * indicates a non-specific (likely IgG related) band.

### TRIB1 loss upon CHX occurs more rapidly than bulk ubiquitin

The mechanisms triggering the loss of TRIB1 protein were then investigated. First, loss of TRIB1 and bulk ubiquitination in the presence of BTZ were compared. Whereas TRIB1 loss in response to CHX was pronounced, loss of coupled Ub was minimal over 5 h (∼23% and ∼ 13% signal loss in HepG2 and HuH-7, respectively), consistent with extensive proteasome inhibition and suggesting that the mechanisms responsible for TRIB1 and bulk ubiquitination removal are distinct (Fig 11).

**Fig 11.**
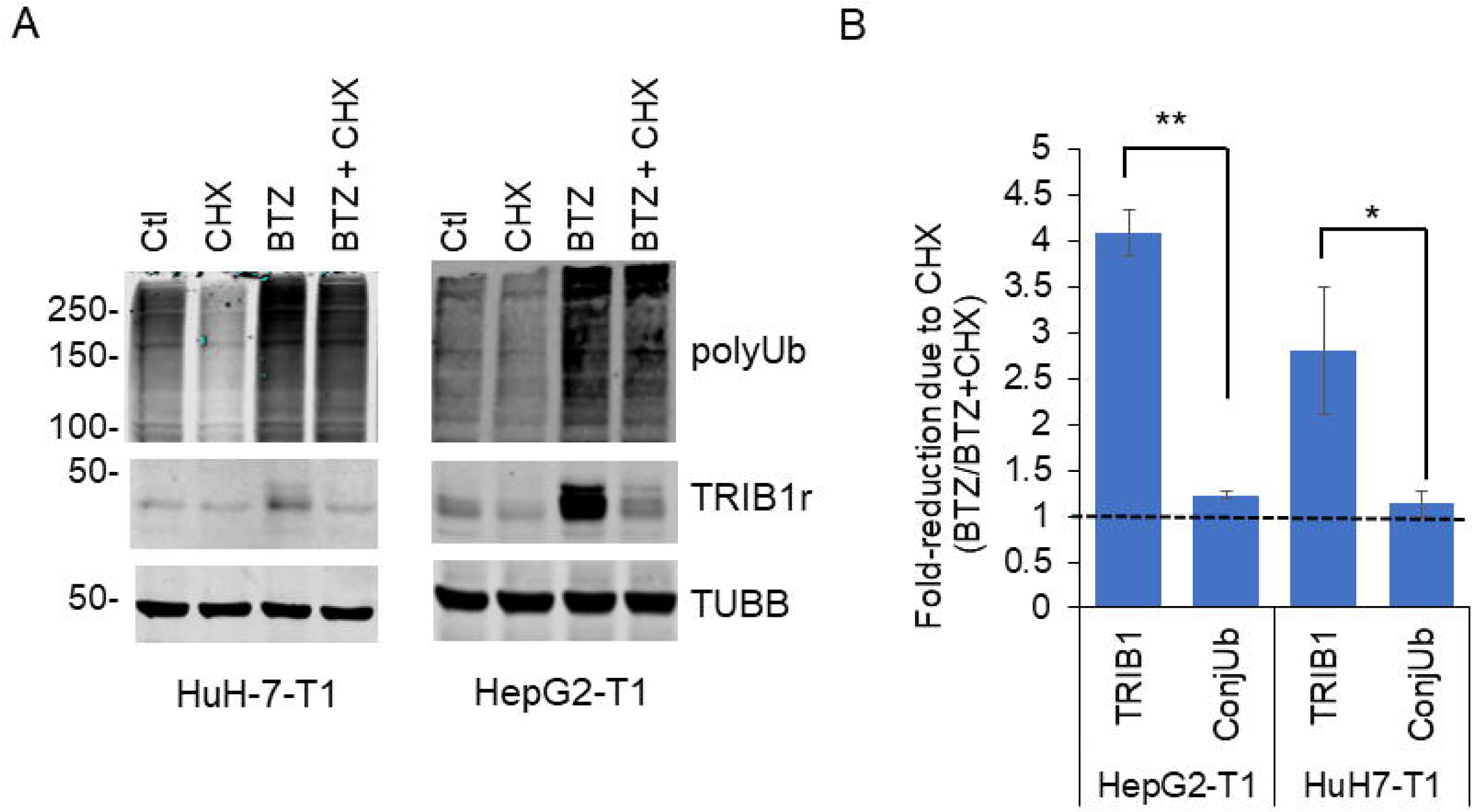
Loss of polyUb in response to CHX is slower compared to TRIB1 in the presence of BTZ. Western blot analysis of HuH7-T1 and HepG2-T1 treated with the indicated drugs for 5 h. A, representative Western blots. Levels of polyUb were quantified over the 100-150 kDa range. B, Relative sensitivities of TRIB1 and polyUb signals to BTZ and CHX after a 5 h treatment. Quantifications of the TRIB1 signals and polyUb (TUBB corrected) in HepG2-T1 and HuH-7-T1 cells following BTZ and/or CHX treatments. Dashed line indicates a no-loss scenario. Average of 3 independent experiments. *, p < 0.05; **, p<0.01; Unpaired T-test, TRIB1 vs ConjUb.

### High dose of Carfilzomib or proteasome inhibitor cocktails do not prevent TRIB1 loss to CHX

The proteasome contains three distinct activities with different sensitivities to PI classes. Indeed, BTZ preferentially targets the chymotrypsin-like and caspase-like activities of the proteasome over the trypsin-like activity. Thus, CHX mediated loss of TRIB1 upon PI treatments may reflect incomplete inhibition of the proteasome, possibly with preferential engagement of the tryptic activity. Carfilzomib is an irreversible, clinically relevant, epoxyketone inhibitor that blocks trypsin-like and chymotrypsin-like activities at high doses *in situ* [40]. Inclusion of Carfilzomib had no impact on TRIB1 loss however (S7A Fig). To further exclude a proteasome contribution, a combination approach was undertaken next. Combined PI use has been shown to increase inhibition potency, possibly reflecting allosteric facilitation [40]. HepG2-T1 cells were incubated in the presence of 3 mechanistically distinct inhibitors: BTZ, LACTA and EPOXO. Combining, all 3 inhibitors did not curb TRIB1 loss however, further minimizing the likelihood of proteasome implication (S7B Fig).

### Proteasome suppression via PSMB3 silencing fails to prevent TRIB1 loss to CHX

A putative contribution of the proteasome machinery to TRIB1 degradation under chemical inhibition was further tested by examining TRIB1 loss to CHX after PSMB3 silencing and PI cocktail; we reasoned that PSMB3 silencing may reveal putative inhibitor resistant residual proteasome activity. As demonstrated earlier (Fig. 9), *PSMB3* silencing was associated with increased bulk protein ubiquitination, indicating impaired proteasome function, yet did not help stabilize TRIB1 in the presence both BTZ and CHX (Fig 12).

**Fig 12.**
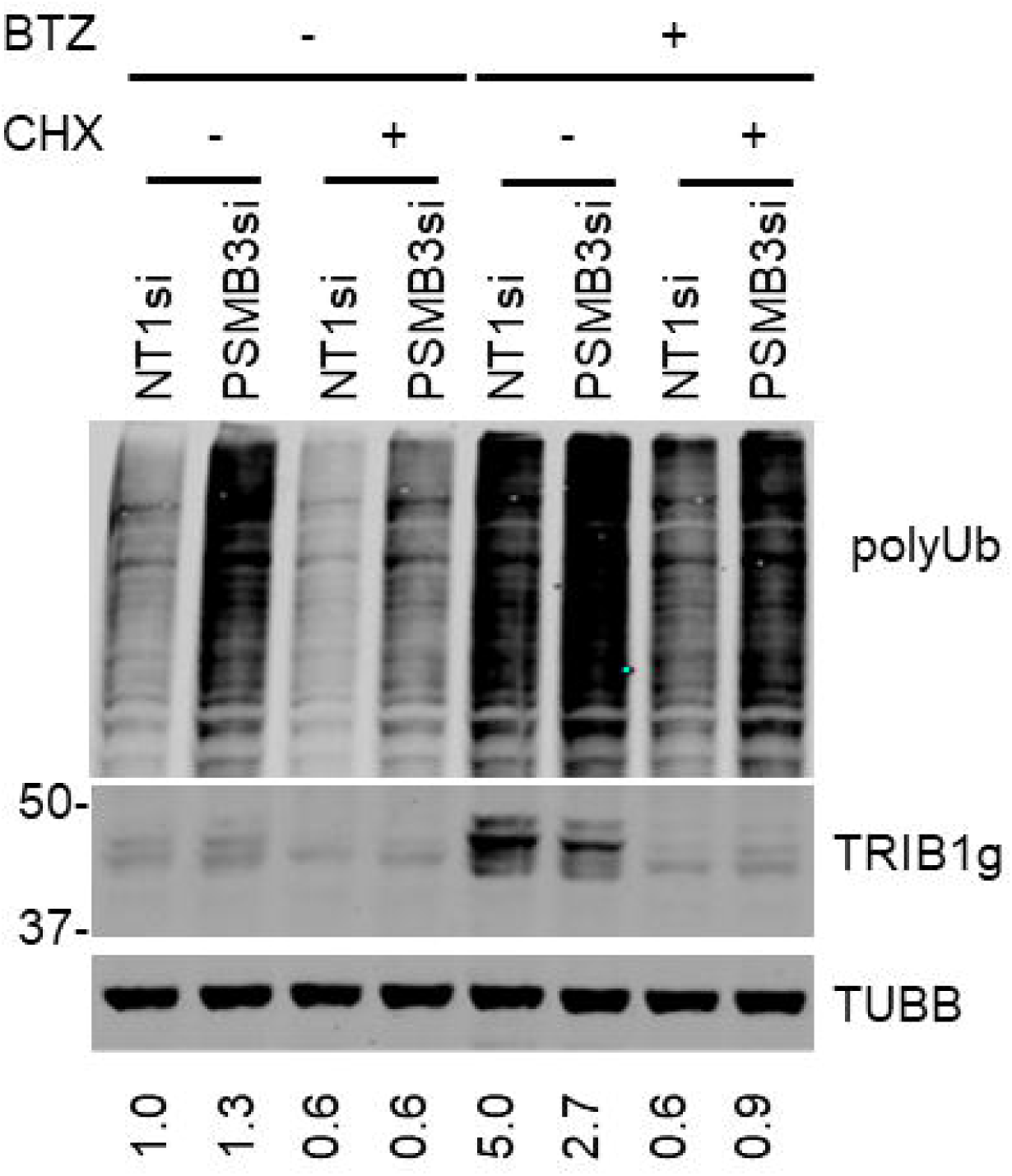
Proteasome-silencing fails to prevent TRIB1 loss to CHX. Western blot analyses of PSMB3-silenced HepG2-T1 cells. HepG2-T1 cells were treated 96 h with siRNA targeting PSMB3 or a siRNA control. During the last 5 h, cells were pre-treated for 5 min with BTZ, followed by the inclusion of CHX for 5 h. For polyUb, region spanning 75 kDa to the top of the gel is displayed. Quantification of the TRIB1g/TUBB signals (normalized to NT1si) is shown.

### COP1 does not regulate TRIB1 protein abundance and turnover

COP1 is an E3 ligase functionally integrated, albeit not exclusively, with TRIB1. It normally resides in both nuclear and cytoplasmic compartments and can shuttle between the 2 compartments [41]. To test whether COP1 plays a role in the loss of TRIB1 in the presence of CHX (under conditions where the proteasome is functional), a TRIB1 construct deficient in COP1 binding, as well as a Wild-type version for direct comparison, were transduced in HepG2 [42]. Both variants were similarly sensitive to CHX, demonstrating TRIB1 instability does not require COP1 interaction (Fig 13).

**Fig 13.**
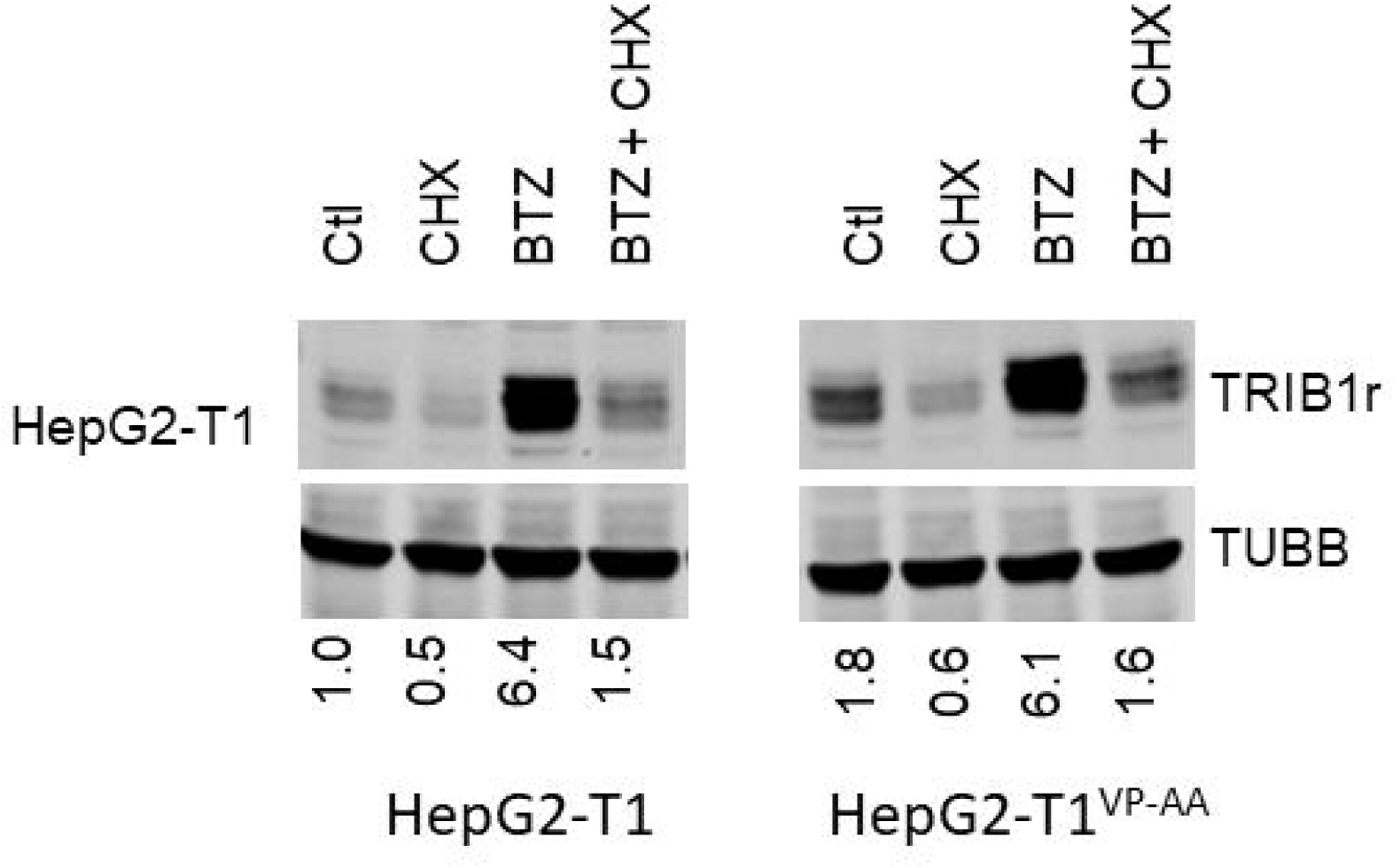
COP1 binding interface is not required for TRIB1 instability. Stable pools of HepG2 cells transduced with either Wt or VP-AA variants were treated with BTZ and/or CHX as indicated for 5 h. Cell lysates were then analyzed by Western blotting using TRIB1 (rabbit) and TUBB antibodies. Data is representative of 2 biologicals. Quantifications are internally controlled to the matching TUBB signal and are expressed relative to the Ctl HepG2-T1.

### Instability of TRIB1 does not require the The Pro-Glu-Ser-Thr (PEST) region or the N-terminal region of TRIB1

The N-terminal region of TRIB2 was implicated in mediating its instability [43]. TRIB1 contains a predicted disordered N-terminus (https://mobidb.bio.unipd.it/Q96RU8; Alphafold) that is enriched in P, E, S and T residues, reminiscent of PEST domains associated with protein instability [44]. To examine whether the PEST-like region played a role in destabilizing TRIB1, we substituted the domain for 2 amphipathic helices of equivalent lengths derived from the more stable GST protein. The substituted TRIB1 was potently upregulated by BTZ but remained highly unstable to CHX, implying that the PEST region is not necessary to confer TRIB1 instability (Fig 14). Similar conclusions were drawn from N-terminal truncations: deletion of either AA 2-51 or AA 2-91 of TRIB1 did not render TRIB1 resistant to CHX (Fig 14). Interestingly, the latter deletion constructs did not show the cluster of bands typical of full-length TRIB1, hinting that this feature is dictated by its N-terminal region and is unrelated to its instability.

**Fig 14.**
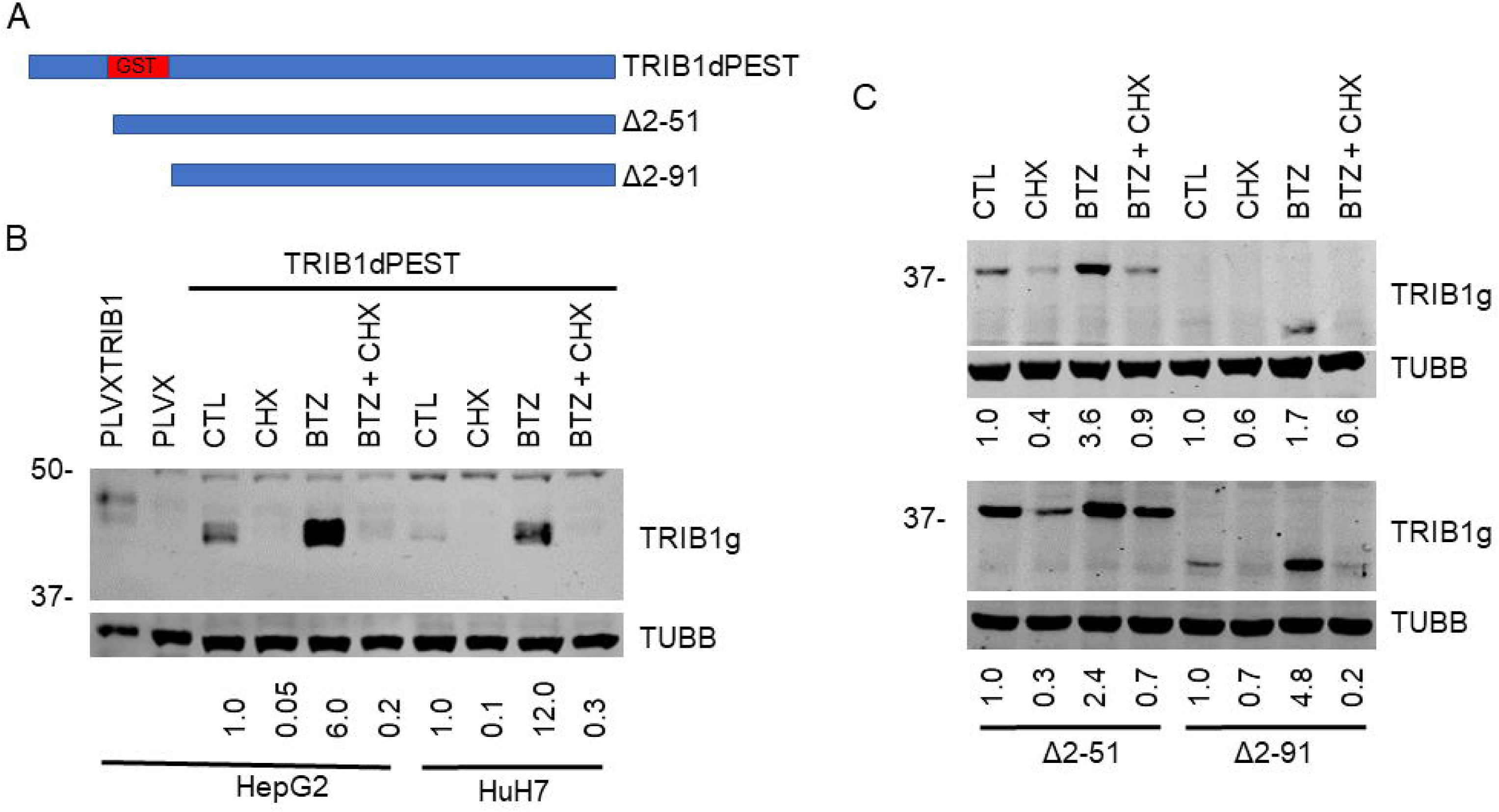
Substitution of TRIB1 PEST region with a GST helical region or N-terminal deletions fail to stabilize TRIB1. A, approximately to-scale schematic of the deletions. B, C, Western blot analysis of stable pools of HepG2 and HuH7 cells expressing TRIB1 with PEST to GST domain substitutions (dPEST) or harboring N-terminal deletions were treated as indicated with CHX and/or BTZ for 5 h. In C, Top, HuH-7-ΔT1; bottom, HepG2-ΔT1 cells. Experiment was repeated twice with similar results.

### Cytosolically retained TRIB1 is unstable

TRIB1 instability could result from regulatory events taking place early during TRIB1 synthesis in the cytosol or after its nuclear import. To test the relative contribution of cytosolic and nuclear processes, we took advantage of a TRIB1 construct wherein the bipartite nuclear localization sequence was replaced PKIA nuclear export signal. We previously showed using transient transfection that TRIB1 was unstable in HeLa cells. To mitigate issues noted previously relating to the greater stability of transiently transfected TRIB1, TRIB1 was transduced by lentiviral delivery. As previously described in non-hepatic cell models, the protein localized exclusively to the cytoplasm (S8 Fig). Inclusion of BTZ resulted in its upregulation, although substituted TRIB1 (TRIB1dNLS) was increased only 2-fold vs ∼4-to 8-fold, for Wt TRIB1 in HepG2 and HuH-7, respectively (Fig 15). Importantly, upon CHX addition, TRIB1 was rapidly lost, in line with degradation processes occurring in the cytosol prior to its nuclear import.

**Fig 15.**
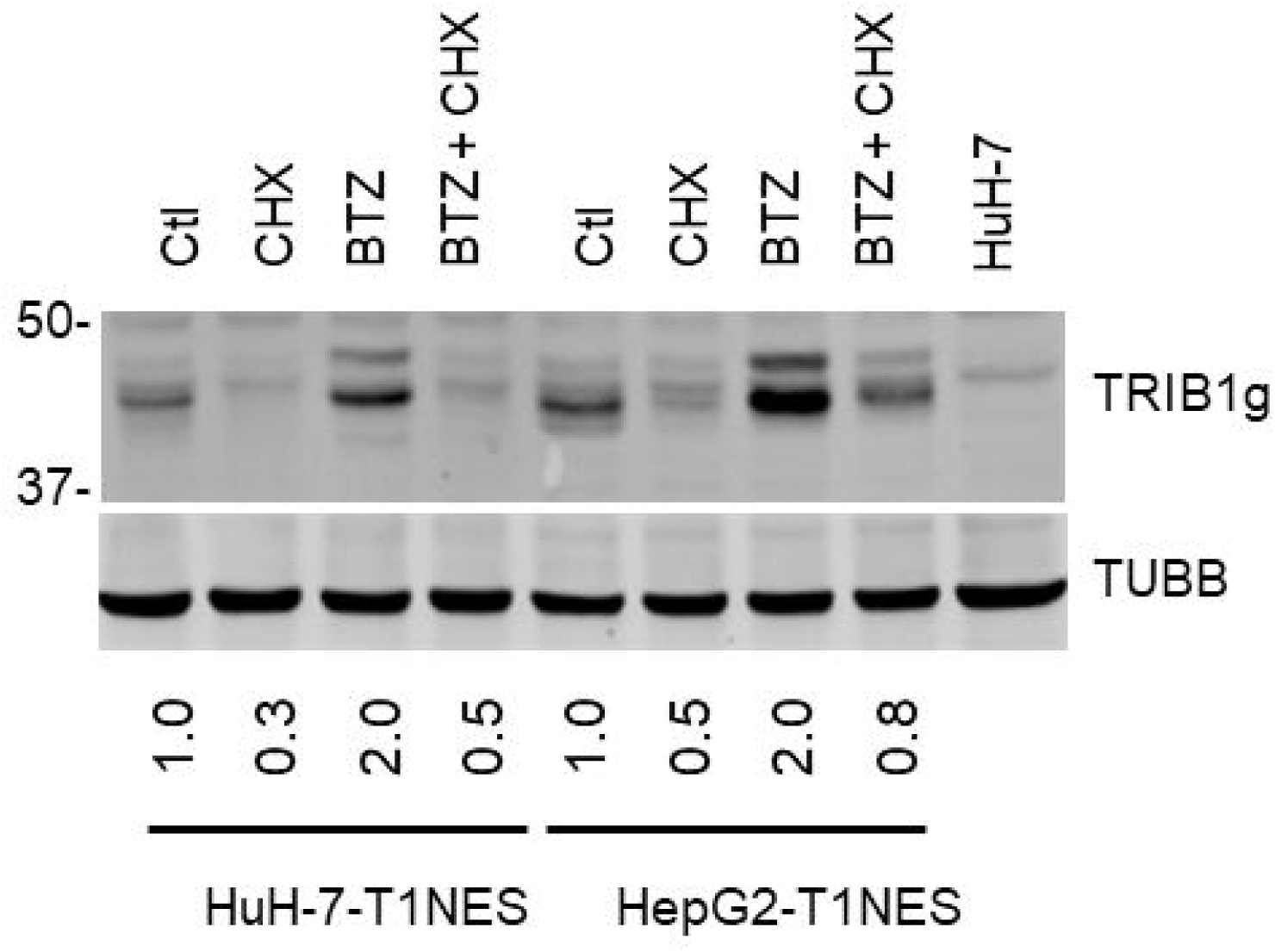
Cytosol-retained TRIB1 is upregulated by BTZ and is sensitive to CHX. Stable pools of TRIB1NES expressing cells generated in HepG2 and HuH-7 cells. Cells were treated for 5 h with BTZ in the presence or absence of CHX. Data are representative of 3 independent experiments.

### TRIB1 proteolysis occurs independently of the lysosomal system and is resistant to protease inhibitors *in vivo*

The lysosomal proteolytic machinery uses its own set of proteases and is largely independent of the proteasome, at least under normal conditions. Proteasome inhibition however was shown previously to activate the lysosomal-autophagy which we hypothesized could be responsible for TRIB1 degradation; this would be consistent with our evidence of cytoplasmic TRIB1 degradation [45]. A possible contribution of the lysosome was tested using CLQ, an inhibitor of lysosome acidification, which is required to activate resident proteases. Prolonged incubation of HepG2-T1 with CLQ, which increased LC3BII, an indicator of lysosome-autophagy inhibition, failed however to prevent recombinant TRIB1 loss (Fig 16). Moreover, basal TRIB1 was not affected by CLQ, which has been previously shown to impair proteasome function (albeit in fibroblasts), further ruling against a contribution of the UPS to TRIB1 degradation [46]

**Fig 16.**
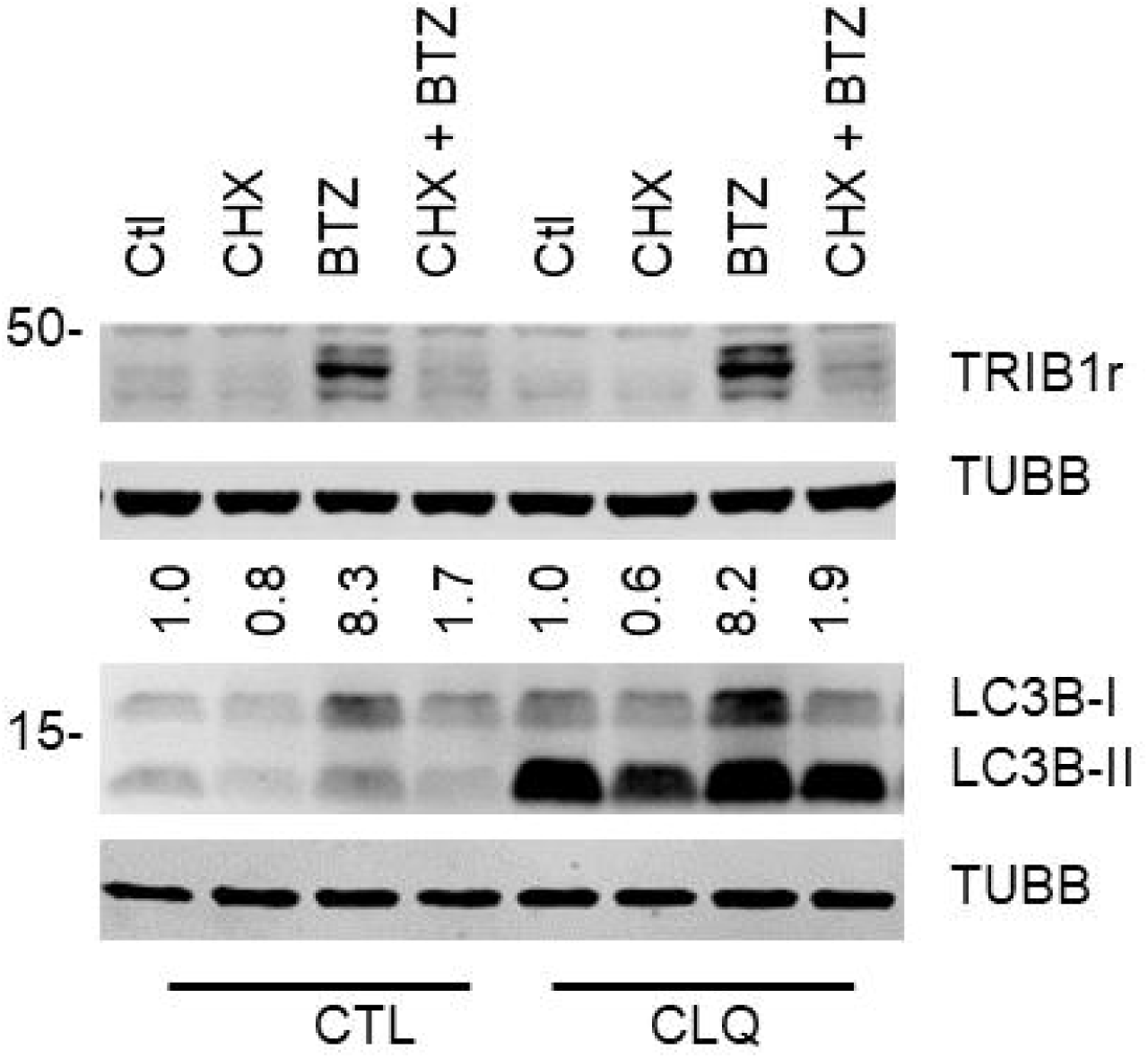
Autophagy inhibition does not impact TRIB1 instability. Western blot analysis of HepG2-T1 cells incubated for 24 h in the presence of CLQ (5 µM) to block autophagy. BTZ and CHX were added during the last 5 h of incubation. LC3B and TRIB1r samples were resolved on 8% and 15% gels, respectively. Experiment was repeated twice with similar results.

Lastly, the contributions of putative proteases were tested by inhibitor screens in HepG2-T1 cells in the presence of BTZ. Broad specificity compounds targeting the major classes of proteases (Ser, Cys and Asp proteases) were selected. Inclusion of cell-permeant protease inhibitors targeting Serine (Pefabloc/AEBSF), Cysteine (E64 and TPCK) and Aspartic acid proteases (Pepstatin A) as well as Calpains (CALPi) and Caspases (ZVAD-FMK) failed to prevent TRIB1 loss resulting from CHX treatment (S9 Fig).

## Discussion

Here we characterized TRIB1 accumulation by proteasome inhibition in hepatocyte models. We show this to be accompanied by increased transcription, as well as instability under proteasome permissive and non-permissive conditions. TRIB1 abundance was strongly increased, largely through transcriptional upregulation but remained unstable in the hepatocyte models examined. This differs from our previous work using non hepatocyte models, wherein we reported a more limited contribution of transcript upregulation (i.e. insensitivity to ACTD) in controlling TRIB1 protein levels following BTZ treatment [29]. We hypothesize that this in part stems from greater TRIB1 mRNA stability, rendering the protein steady state levels less sensitive to ACTD. Importantly TRIB1 remained highly sensitive to CHX in the presence of BTZ, indicating that proteasome-independent protein instability is conserved in all models examined to date.

A possible confounder is the ability of cells to detoxify and/or rapidly export non-covalent inhibitors (e.g BTZ). However, the incubation conditions (relatively short, continuous and in the presence of high concentrations), combined with very slow off rates (for BTZ at least [47]) should minimize these possibilities. Moreover, stability of the poly-Ub signal observed in the presence of PIs indicates that proteasome inhibition is persistent. Although it cannot be totally excluded that low, residual activity is sufficient to degrade TRIB1, a lack of proteasome involvement in TRIB1 degradation is further supported by the absence of identifiable Ub-TRIB1 complexes, with the caveat that Ub-TRIB1 abundance may be too low to detect. Absence of Ub signal does not exclude a proteasome contribution however: the 20S proteasome, which lacks the gatekeeper 19S regulatory particle responsible for ubiquitin recognition, can promote degradation of ubiquitinated targets [48, 49].

The proteasome is responsible for the vast majority of protein degradation, including most short lived proteins [50]. Co-translational degradation via the UPS is widespread and contributes to protein synthesis quality control [51–54]. It has been estimated that 12-15% of nascent polypeptides are ubiquitinated in HEK293T [51]. In that case, in contrast to TRIB1, de novo co-translated proteins are ubiquitinated and can be readily stabilized by proteasome inhibition. Non UPS proteolytic pathways, sometimes detectable only in the absence of functional proteasome, have been described, including Calpains, TPP2 and others (e.g. [55–59]). According to the MEROPS database, the human proteome contains 703 putative and validated proteases, most of which await elucidation of biological role and might function in more specialized forms of intracellular proteolysis.

In contrast to TRIB1, TRIB2 has been shown to be subjected to ubiquitination via smuf1 and/or bTRCP, and possibly proteasome-mediated degradation, although this remains to be formally tested via an inhibitor approach [43, 60]. Thus, despite their shared origins, divergent regulatory mechanisms may have co-evolved to regulate individual tribbles. Indeed, while TRIB2 instability to CHX treatment (when overexpressed in HE293T) was largely abrogated by removal of its disordered N-terminal 50 AA, TRIB1 deletions spanning the N-terminus as well as substitutions of AA 31 to 88 (NLS + PEST regions) were insufficient to stabilize TRIB1. In addition, TRIB2 can be destabilized pharmacologically via an interaction between its pseudo kinase domain and a family of EGFR inhibitors [61]. As all tribbles proteins harbor somewhat conserved pseudo kinase domain, this domain may render TRIB1 unstable. Instability could be mediated via post-translation modifications (PTMs). For instance, for both cyclin D1 and AKT, phosphorylation impairs (proteasome-mediated) degradation [54, 62]. Relatively little is known about TRIB1 PTMs. TRIB1 migration on SDS-PAGE, characterized by several closely migrating bands, is certainly consistent with low molecular weight covalent modifications (e.g., phosphorylation). Interestingly, an analogous pattern was observed with terminal the PEST- and NLS-substituted TRIB1 (AA 53-88 and AA 30-51, respectively) construct but not with the Δ2-51 N-terminal deletion, implying that this pattern requires AA 2-29 of TRIB1.

While we present evidence excluding possible degradative pathways, the how and why of TRIB1 instability remain unknown. Recognition subunits of E3 ligase, like TRIB1, tend to be rapidly turned over, suggesting that instability is functionally important [16]. Interestingly, TRIB1 instability appears independent of its interaction with COP1, as suggested by the propensity of the binding deficient variant to remain unstable. Rapid TRIB1 turn over might stem from quality control events, perhaps to address a propensity of TRIB1 to misfold. Alternatively, TRIB1 may be specifically turned over to ensure a timely adaptation to changing cellular conditions. Given the role of TRIB1 in cell growth and proliferation, one could envision interactors either stabilizing or destabilizing the emerging TRIB1 to match nutrient abundance. Future work will explore these possibilities.

## Acknowledgments

This work was funded by a Canadian Institutes for Health Research Foundation grant (FRN:154308; RM).

## Supporting information captions

S1 Fig. TRIB1 mRNA increases in response to proteasome inhibitors in HuH-7 and HepG2 cells. TRIB1 mRNA abundance (normalized to PPIA level) was measured following 5 h regimen with the indicated drugs. Values are expressed relative to the vehicle (1 % DMSO). Data represent the average of 3 experiments ± S.D. All changes from vehicle (except for LACTA 2 µM; HepG2) were statistically significant (p<0.05)

S2 Fig. Endogenous TRIB1 signal is specific, nuclear and is detectable in HepG2 cells following a 5 h BTZ treatment. HepG2 cells were seeded on coverslips and treated as indicated with BTZ (5 µM) and/or CHX (10 µg/ml). Cells were then fixed and permeabilized prior to incubations with either a TRIB1 antibody (1:500, +) or no antibody (-) for 1 h. Cells were then washed and incubated for 1 h with a Donkey anti-goat (DAG633) secondary Ab (1:2000) and counterstained with Hoechst. Images were smoothed with a Gaussian filter.

S3 Fig. Increased TRIB1g reactive nuclear signal in HuH-7 treated for 18 h with BTZ 5 µM. A, wide field (60 X). CARS2 is included as a cytoplasmic marker. B, magnification of yellow boxed region shown on left, smoothed with Gauss filter processing. TRIB1g (peudocolored red), Hoechst’s (pseudocolored lime green).

S4 Fig. BTZ induces changes consistent with the establishment of an ER stress response in HepG2 and HuH-7 cells. Treatment with BTZ (5 h) increases BiP and CHOP mRNA levels. Values were first normalized to PPIA and are expressed relative to vehicle (1% DMSO). Paired Student t-test, BTZ vs vehicle. *, P<0.05; **; P<0.01

S5 Fig. Increased recTRIB1 upon proteasome treatment. HuH-7 and HepG2 cells stably expressing TRIB1 were incubated with vehicle (Ctl), LACTA, BTZ or EPOXO at the indicated concentrations for 5 h. A, protein abundance assessed by Western blot; B, mRNA abundance as measured by qRT-PCR Data representative of 2 experiments.

S6 Fig. PSMB3 suppression leads to increased TRIB1 mRNA abundance in HepG2-T1 but not HuH7-T1 cells. PSMB3 was suppressed for 72 h and mRNA abundance was measured by qRT-PCR. mRNA levels were corrected for PPIA and are expressed relative to a non-specific control (NT).

S7 Fig. High doses carfilzomib or a proteasome inhibitor cocktail does not prevent TRIB1 loss to CHX in HepG2-T1 cells. Western blot analyses of HepG2-T1 cells treated with proteasome inhibitors. In A, HepG2-T1 cells were treated with the indicated carfilzomib, either alone or in the presence of CHX for 5 h; carfilzomib was added 5 min prior to CHX (10 µg/ml). In B, cells were treated for 5 h with a 5 min preincubation with BTZ (B), vehicle (Ctl, 1% DMSO) or a combination of BTZ (B, 5 µM), Lactacystin (L, 10 µm) and Epoxomicin (E, 2 µm) and/or with 10 µg/ml cycloheximide (CHX), where indicated. Experiments were repeated twice with similar results. Quantifications are the TUBB-corrected TRIB1 signal and are normalized to the Ctl value (B) or the matching CHX-free value (A). Experiments were repeated twice, with similar results.

S8 Fig. TRIB1NES expression in transduced HuH-7 and HepG2. HuH-7 and HepG2 cells stably transduced with TRIB1NES were treated for 5 h with BTZ. Fixed cells were permeabilized and incubated for 16 h with TRIB1g antibody. T1g was detected with an Alexa 633 donkey anti-goat Ab. Nuclei were counterstained with Hoechst.

S9 Fig. Preincubation with protease inhibitors cannot prevent TRIB1 loss in the presence of CHX and BTZ. HepG2-T1 cells were treated with BTZ in the presence of PEFABLOC (PEFA, 200 µM), Pepstatin A (PEPA, 1 µM), E64 (2 µM), calpain inhibitor 1 (CALPi, 20 µM), ZVAD-FMK (ZVAD, 1 µM) and TPCK (25 µM). Protease inhibitors were added concurrently with BTZ and 5 min prior to the addition of CHX. Data are representative of 3 experiments.

