## Supplementary figures and images for "Regulation of TRIB1 abundance in hepatoma models"

### Supplemental figures

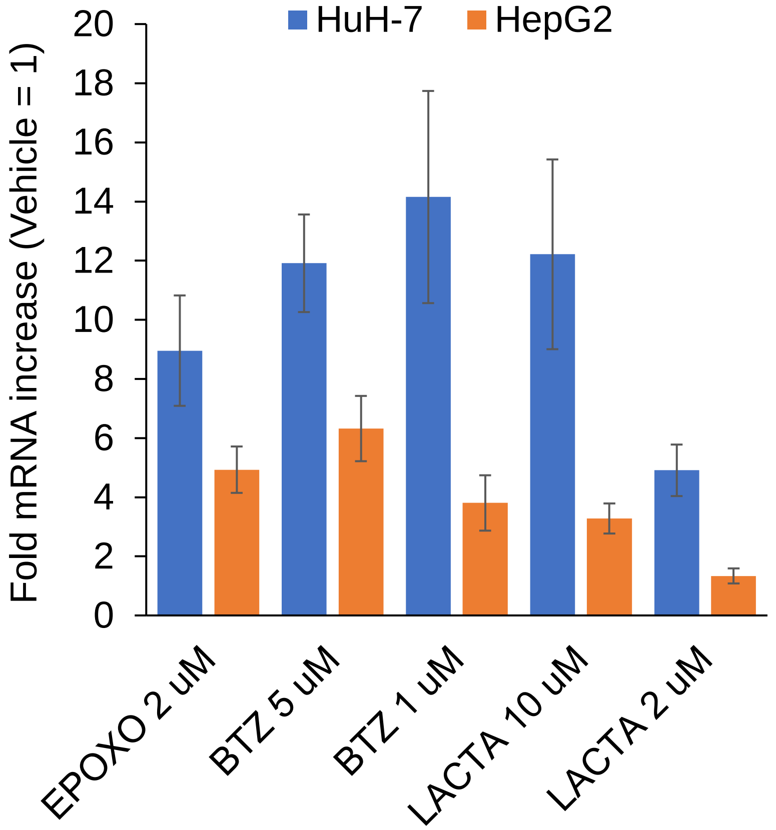

S2 Fig

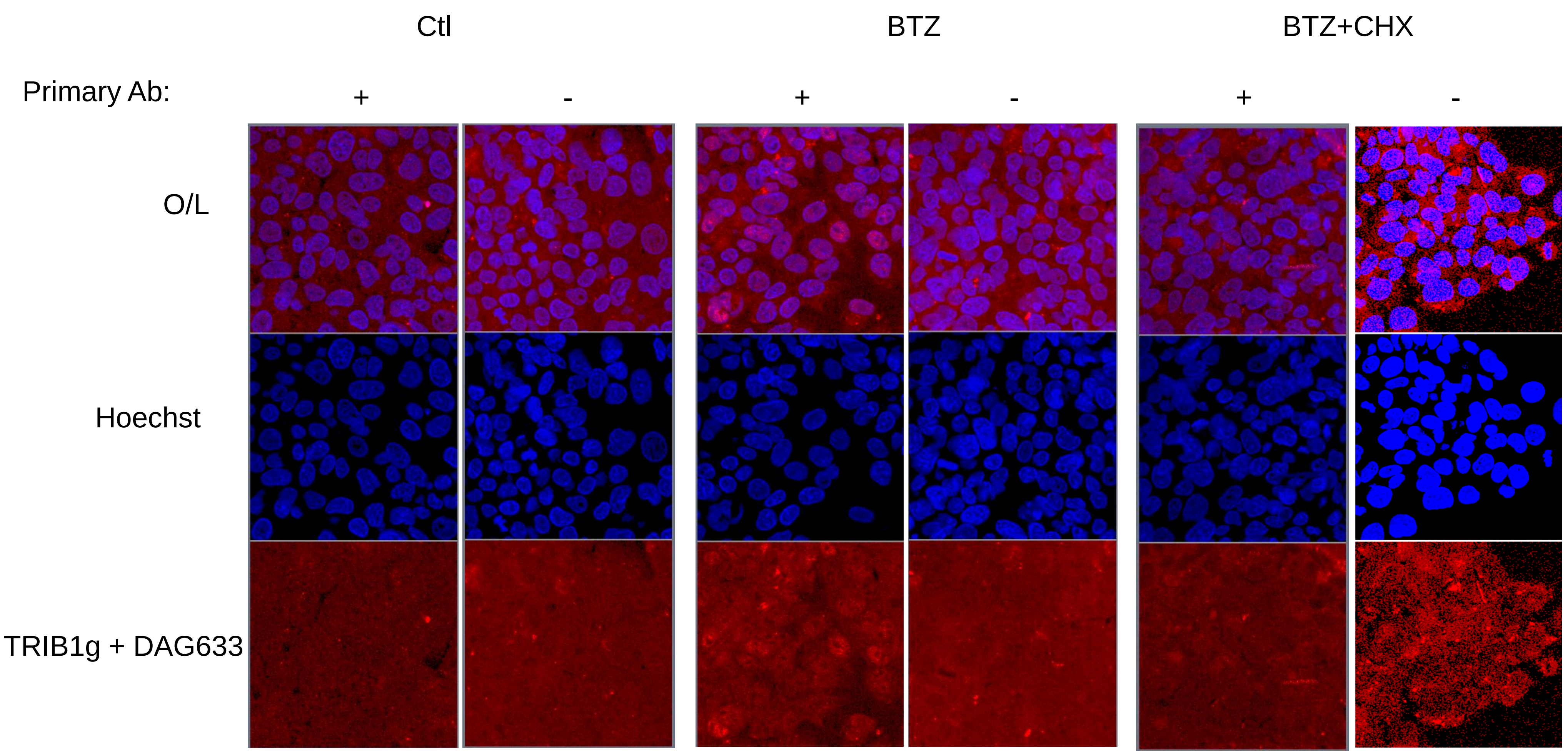

#### S3 Fig

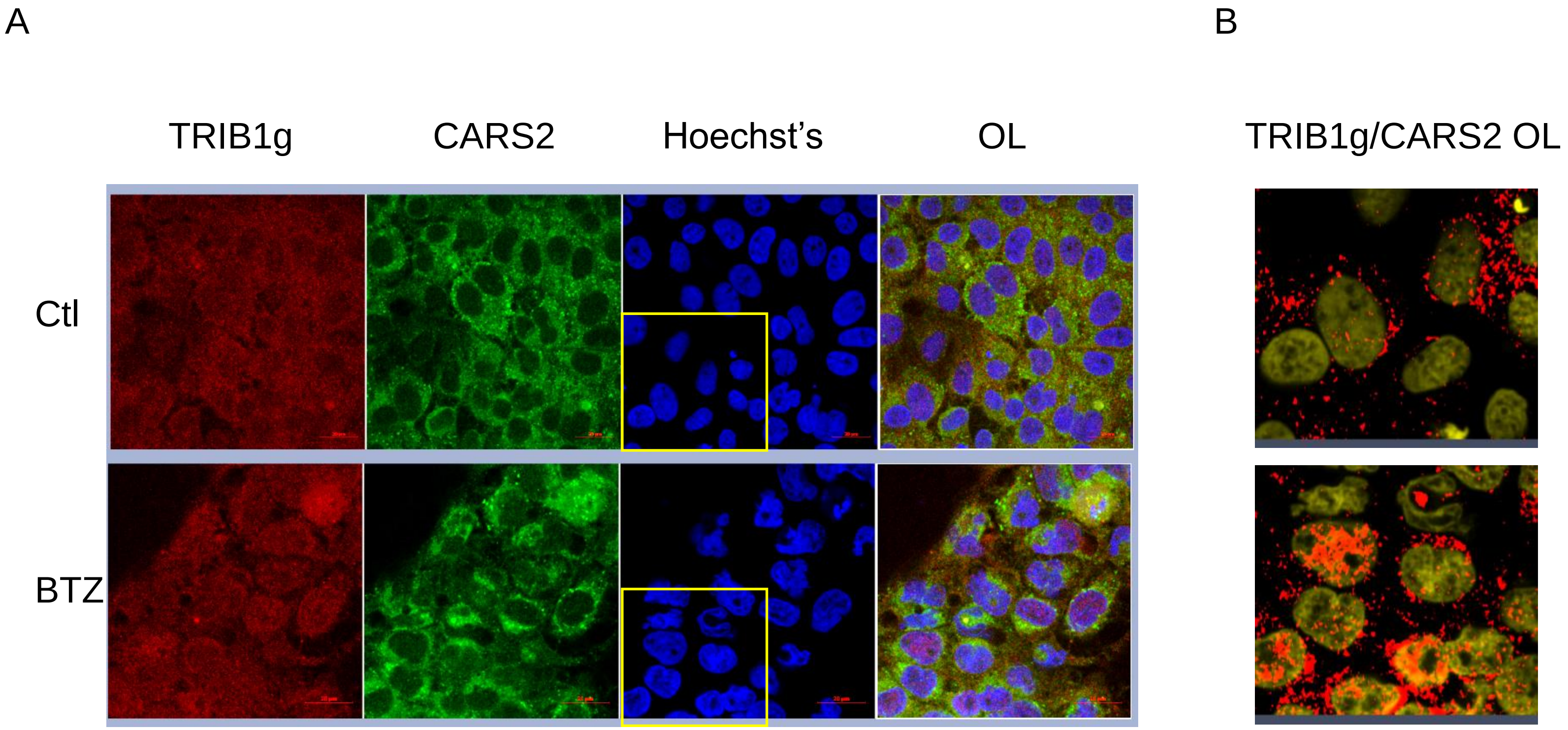

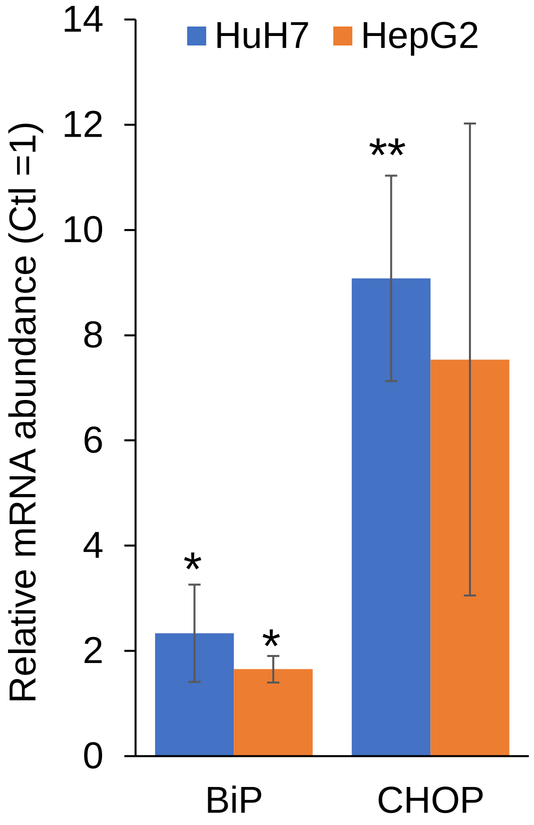

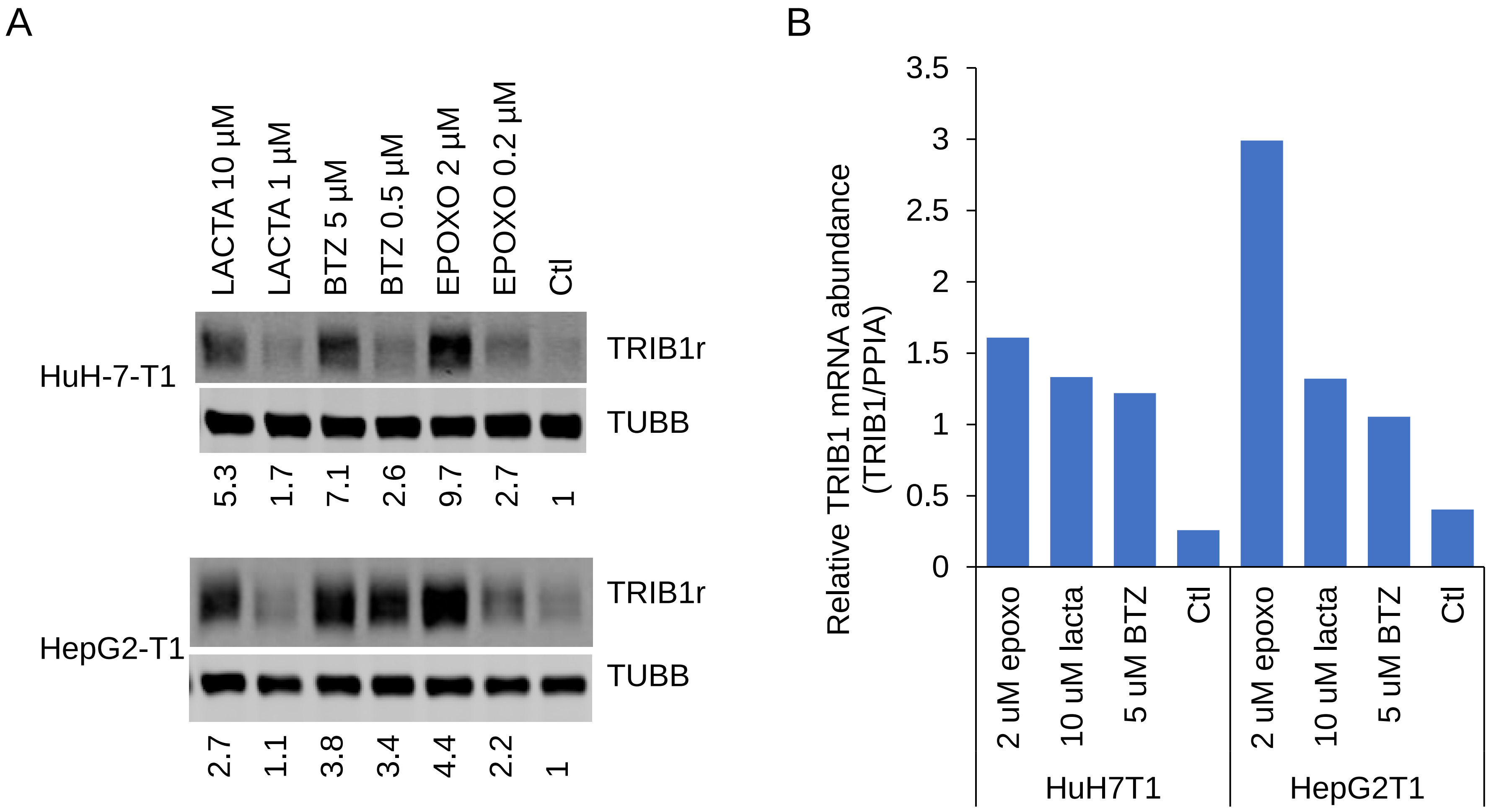

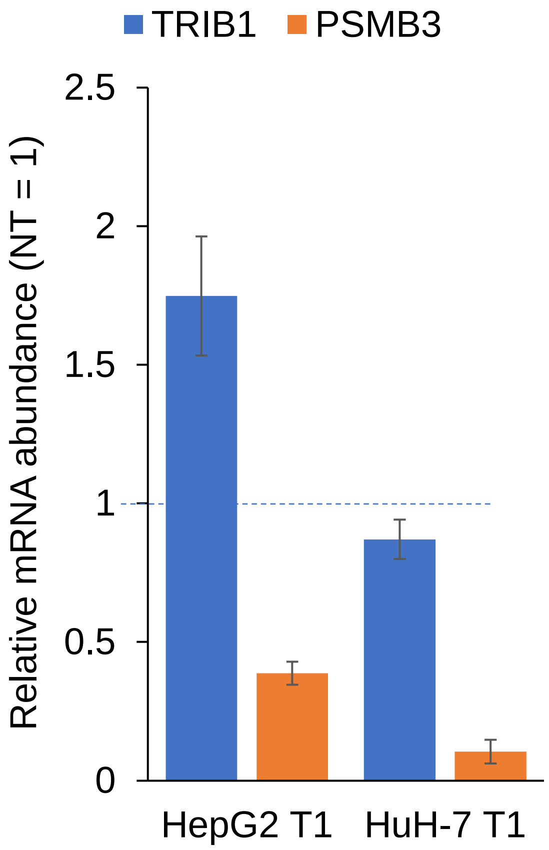

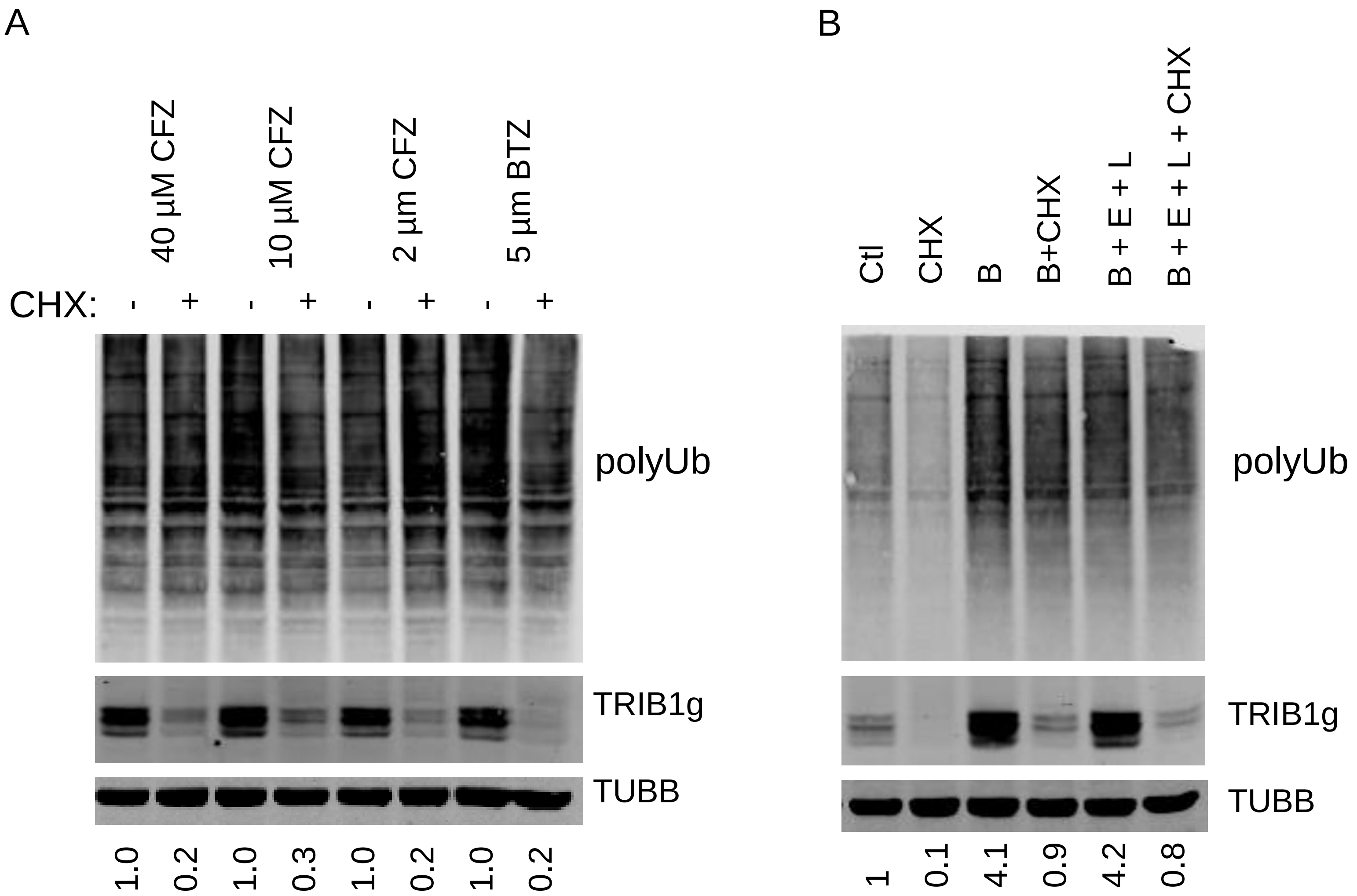

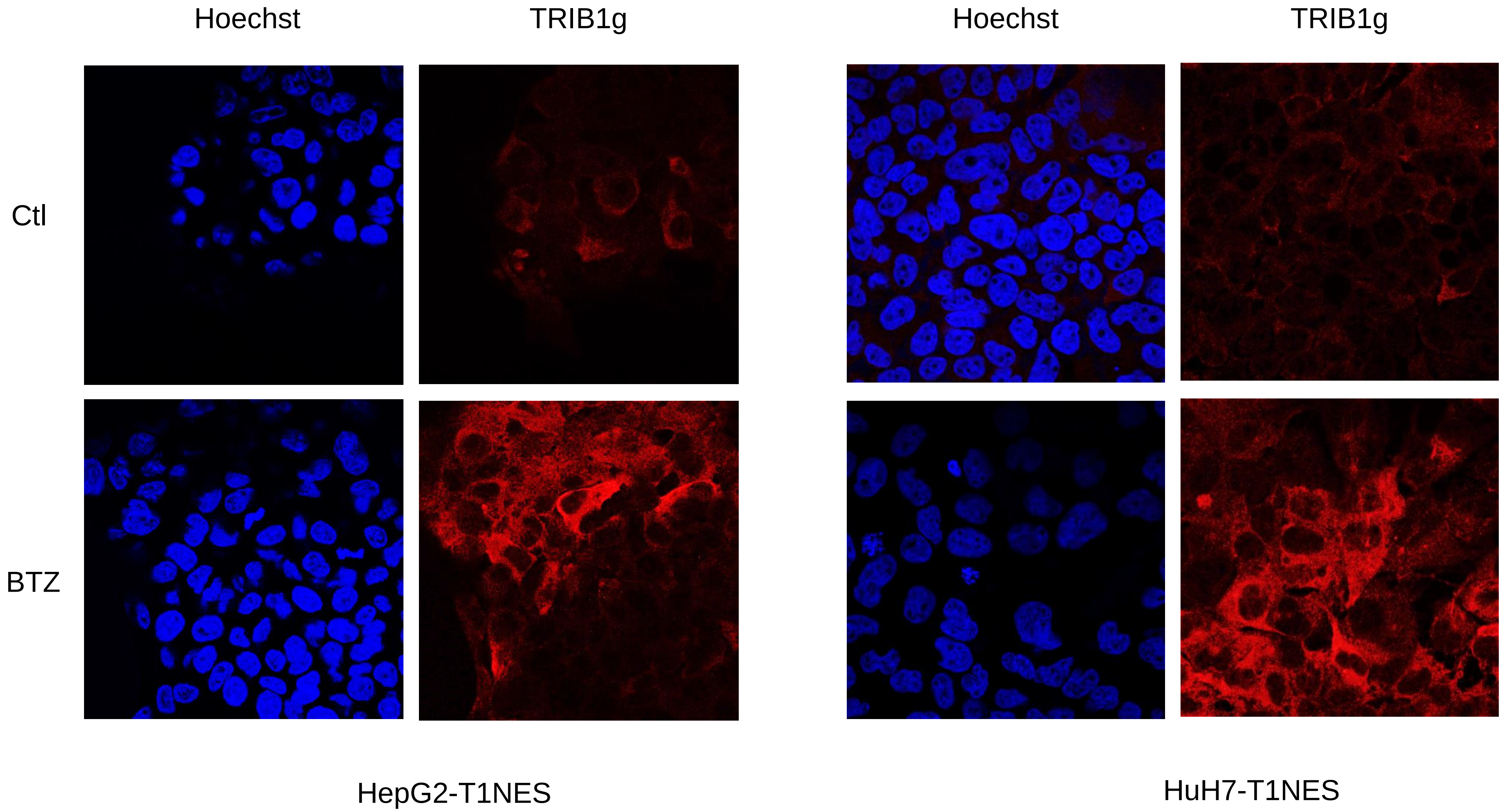

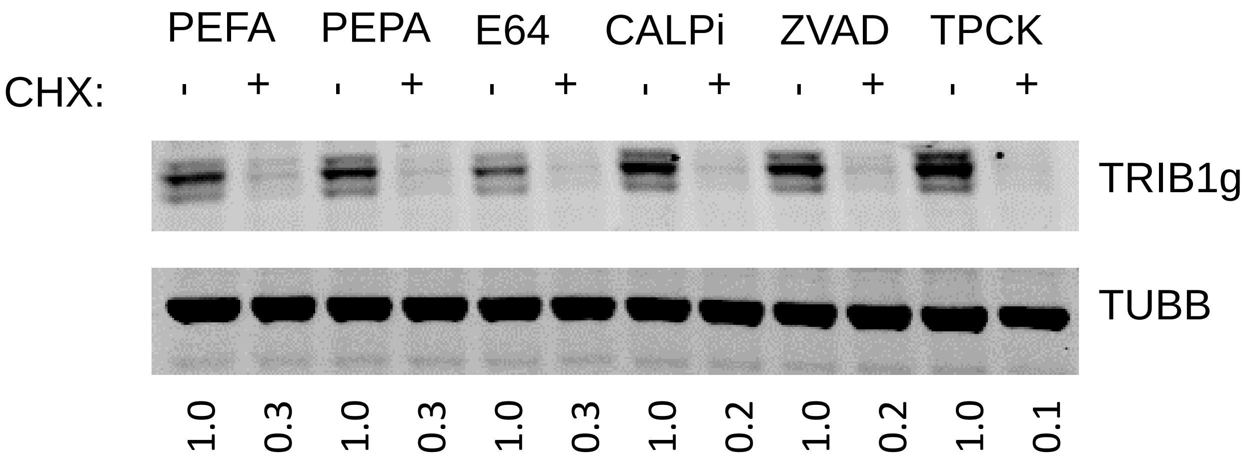
